# Ephemeral states in protein folding under force captured with a novel magnetic tweezers design

**DOI:** 10.1101/310060

**Authors:** Rafael Tapia-Rojo, Edward C. Eckels, Julio M. Fernandez

**Affiliations:** Department of Biological Sciences, Columbia University, New York, NY 10027, USA

## Abstract

Magnetic tape heads are ubiquitously used to read and record on magnetic tapes in technologies as diverse as old VHS tapes, modern hard drive disks, or magnetic bands on credit cards. Their design highlights the ability to convert electric signals into fluctuations of the magnetic field at very high frequencies, which is essential for the high density storage demanded nowadays. Here, we twist this conventional use of tape heads to implement one in a new magnetic tweezers design, which offers the unique capability of changing the force with a bandwidth of *~* 10 kHz. We calibrate our instrument by developing an analytical expression that predicts the magnetic force acting on a superparamagnetic bead based on the Karlqvist approximation of the magnetic field created by a tape head. This theory is validated by measuring the force dependence of protein L unfolding/folding step sizes, and the folding properties of the R3 talin domain. We demonstrate the potential of our instrument by carrying out millisecond-long quenches to capture the formation of the ephemeral molten globule state in protein L, which has never been observed before. Our instrument provides for the first time the capability of interrogating individual molecules under fast-changing forces with a control and resolution below a fraction of a pN, opening a range of novel force spectroscopy protocols to study protein dynamics under force.

---

Magnetic head recording systems have been perfected over decades, resulting in a deep understanding of the physical features of magnetic tape heads, which evolved to create strong magnetic fields that can change very rapidly in time [1, 2]. Hence, it becomes natural that we explore the use of this technology in its application to force spectroscopy. Magnetic tweezers force spectroscopy uses magnetic field gradients to apply pulling forces on biomolecules tethered to superparamagnetic beads [3–9]. Due to the extreme compliance of the magnetic trap, magnetic tweezers offer intrinsic force-clamp conditions and an exquisite control of the pulling force, which, combined with HaloTag covalent chemistry, provides an inherent stability, and gives access to long timescales, of several hours or even days [10–13]. However, in standard magnetic tweezers instruments, the force change is limited by the mechanical movement of the pair of magnets, which can take up to 100 ms, and impedes to capture early molecular events occurring upon fast force quenches, or to apply force protocols where the force changes rapidly, motivating the need to implement fast-changing magnetic fields.

In this article, we present magnetic tape head tweezers, a novel force spectrometer capable of changing the force on a micro-second timescale, while maintaining an impeccable control of it. Thanks to the Karlqvist description of the magnetic field created by a tape head [14], we provide a full analytic description of the pulling force, which allows us to calibrate our instrument over a wide range of forces. We further confirm this approach by using the extremely force-sensitive talin folding dynamics to validate the force dependency of the instrument. Finally, we demonstrate the potential of our setup by directly observing the folding mechanism of protein L at very short timescales. Thanks to the ability to carry out ultra-fast force quenches to well-defined forces, we capture the molten globule state of protein L, which forms in few milliseconds.

## RESULTS

### Calibration of the magnetic tape head tweezers using protein L step sizes

Magnetic tape head tweezers follow the same scheme as conventional magnetic tweezers [6], but substituting the pair of voice coil-mounted magnets by a micro-positioned magnetic tape head (Brush Industries, 902836), identical to those employed in tape recorders or credit card readers (Fig. 1) [15]. A magnetic head consists of a toroidal core of magnetic material with large permeability [1]. The core has a narrow gap (25 *µ*m), over which a strong magnetic field is generated when an electric current is supplied to a coil wrapped around. Compared to other electromagnetic tweezers designs [4, 16, 17], our setup highlights the ability to change the force with a bandwidth of several kHz (*∼* 10 kHz with our implementation, see SI). Just as tape heads are designed to convert electric signals into a magnetic record on a tape, here we employ it to convert electric signals into force signals, applied to tethered molecules (see Fig. 1B).

**FIG. 1.**
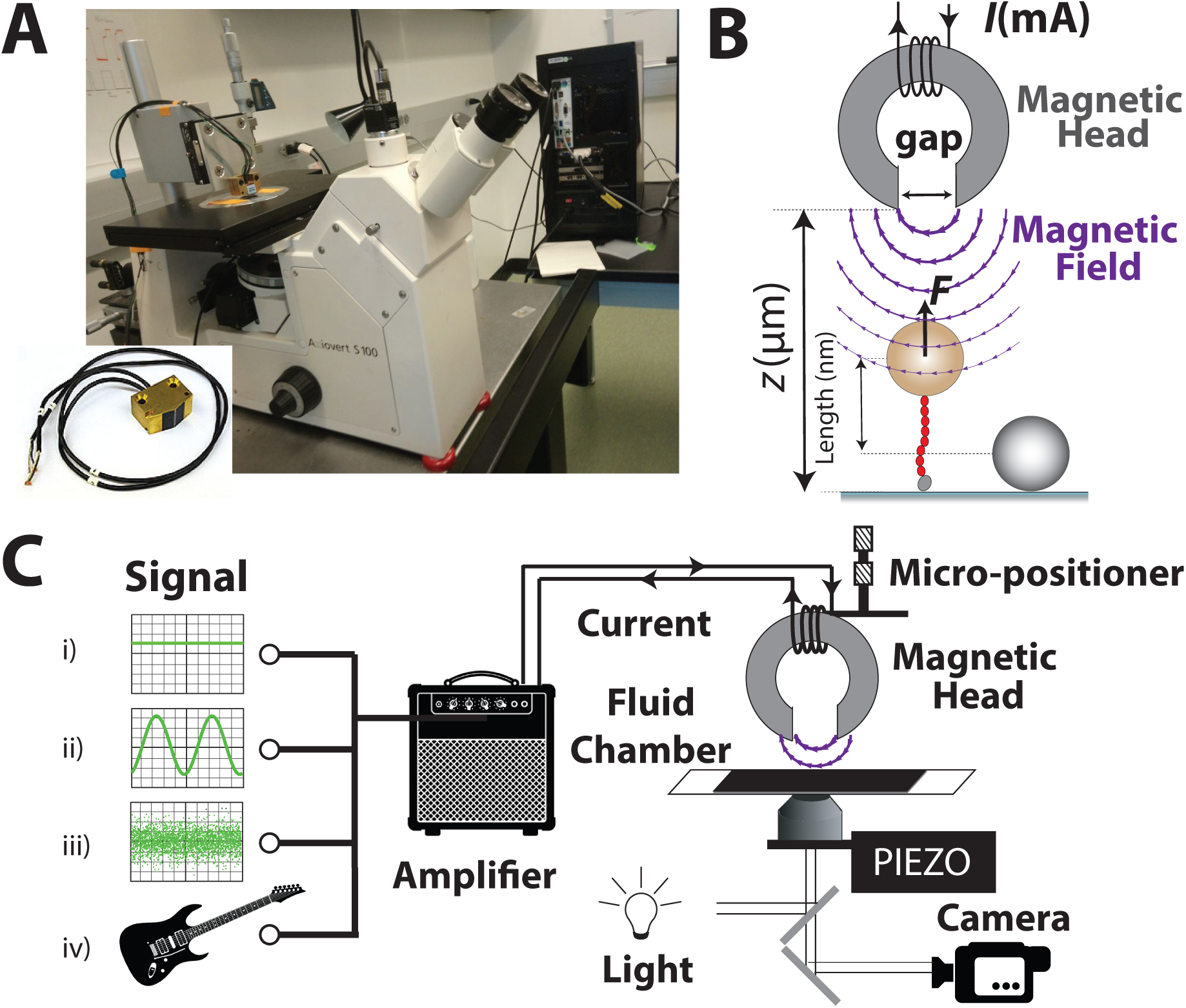
Magnetic tape head tweezers setup. (A) Photograph of the magnetic tape head tweezers setup, highlighting the magnetic tape head. (B) Schematics of the magnetic tape head tweezers experiment. (C) Schematics of the magnetic tape head tweezers setup. Electric current signals of any shape are converted to force signals, which are directly applied on tethered proteins.

In magnetic tweezers, the applied force is determined by using a calibration curve that relates the pulling force with the vertical distance between the magnet and the bead [4–6]. This *magnet law* is specific for the geometry and strength of the magnets and the specifications of the magnetic beads. In our case, the pulling force depends, not only on the distance between the magnetic head and the bead, but also on the supplied current. Following the strategy previously developed [6], we generate a *current law* by relating the pulling force (*F*) with the distance from the head to the bead (*z*) and the electric current (*I*) through the unfolding/folding step sizes of protein L, which scale with the force following standard polymer physics models such as the freely-jointed chain (FJC) model [18]. However, in order to address this problem, we need a description of the dependence of the pulling force with *z* and *I*. In conventional magnetic tweezers, an empirical exponential dependence between force and distance is assumed [6, 19]. However, here we can rely on an analytical relation based on the Karlqvist description of the magnetic field created by a tape head (see SI) [1, 14].

Under an external magnetic field 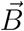, the force acting on a superparamagnetic bead can be described as 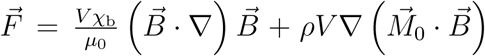 [20], where *ρ* is the density of the bead (kg/m^3^), *V* its volume (m^3^), 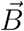 the applied magnetic field (T), *χ*_b_ the magnetic susceptibility of the bead (dimensionless), and *µ*_0_ = 4*π* × 10^−7^ Tm/A [21–23]. Despite using superparamagnetic beads, we include an initial magnetization term 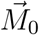, which has been demonstrated to be necessary when working with commercial beads and at weak fields (see [21] and SI Appendix for a discussion on this approximation). The magnetic force on a superparamagnetic bead is often described in the literature only with the first term. However, in actual superparamagnetic beads 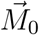 can be comparable to the magnetization induced by weak fields (see [21] and SI).

The lateral *F*_x_ and vertical *F*_z_ components of the magnetic force acting on the bead are obtained by using Karlqvist description of the magnetic field on the force acting on the bead (see SI Appendix for explicit derivation). In practice, we work over the gap region (*−g*/2 ≤ *x* ≤ *g*/2) where the magnetic field is constant and the lateral component vanishes *F*_x_ = 0. Thus, the vertical component evaluated at *x* ≈ 0 can be written as (dropping the z subscript):

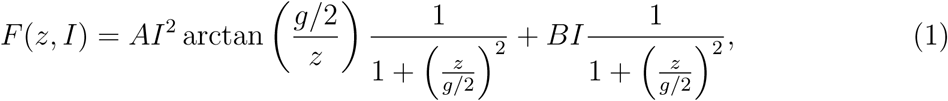

where *A* and *B* are constants built with the physical parameters of our problem (see SI Appendix). Our calibration strategy hinges on determining *A* and *B* by relating the extension ∆*L* of the folding/unfolding steps of protein L—used as a molecular ruler—with *F* through the FJC model, 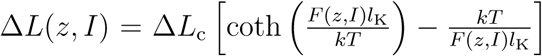, where *F* (*z, I*) takes the shape of Eq. 1, ∆*L*_c_ is the contour length increment, *l*_K_ the Kuhn length (∆*L*_c_ = 16.3 nm and *l*_K_ = 1.1 nm for protein L [10]), and *kT* = 4.11 pNnm. These relation could have been equally derived using a worm-like chain description of the step sizes. However, the FJC model has an explicit ∆*L*(*F*) dependence that makes it analytically more convenient.

We carry out experiments on protein L octamers using the M-270 2.8 *µ*m beads as described previously (see [6] and methods), working at different values of *z* and *I*, and measuring the average step size ∆*L*(*z, I*). Figure 2A shows a typical magnetic tape head tweezers trajectory for a protein L octamer. First, working at *z* = 275 *µ*m, we unfold the octamer with a pulse of 800 mA, and the unfolding of each domain appears as a step-wise increase in the extension of 14.3 nm. Next, we decrease the current to 225 mA, and the polyprotein collapses, showing unfolding/refolding dynamics with steps of 7.9 nm. Finally, we unfold the remaining domains at 665 mA, and move the head away from the bead to *z* = 550 *µ*m, which results in a decrease of the pulling force, as the protein folds to a state where all eight domains are folded on average, showing steps of 5.6 nm. For each (*z, I*) condition, we determine the step sizes as the distance between two consecutive length histograms, calculated over the smoothed trajectory (black trace).

**FIG. 2.**
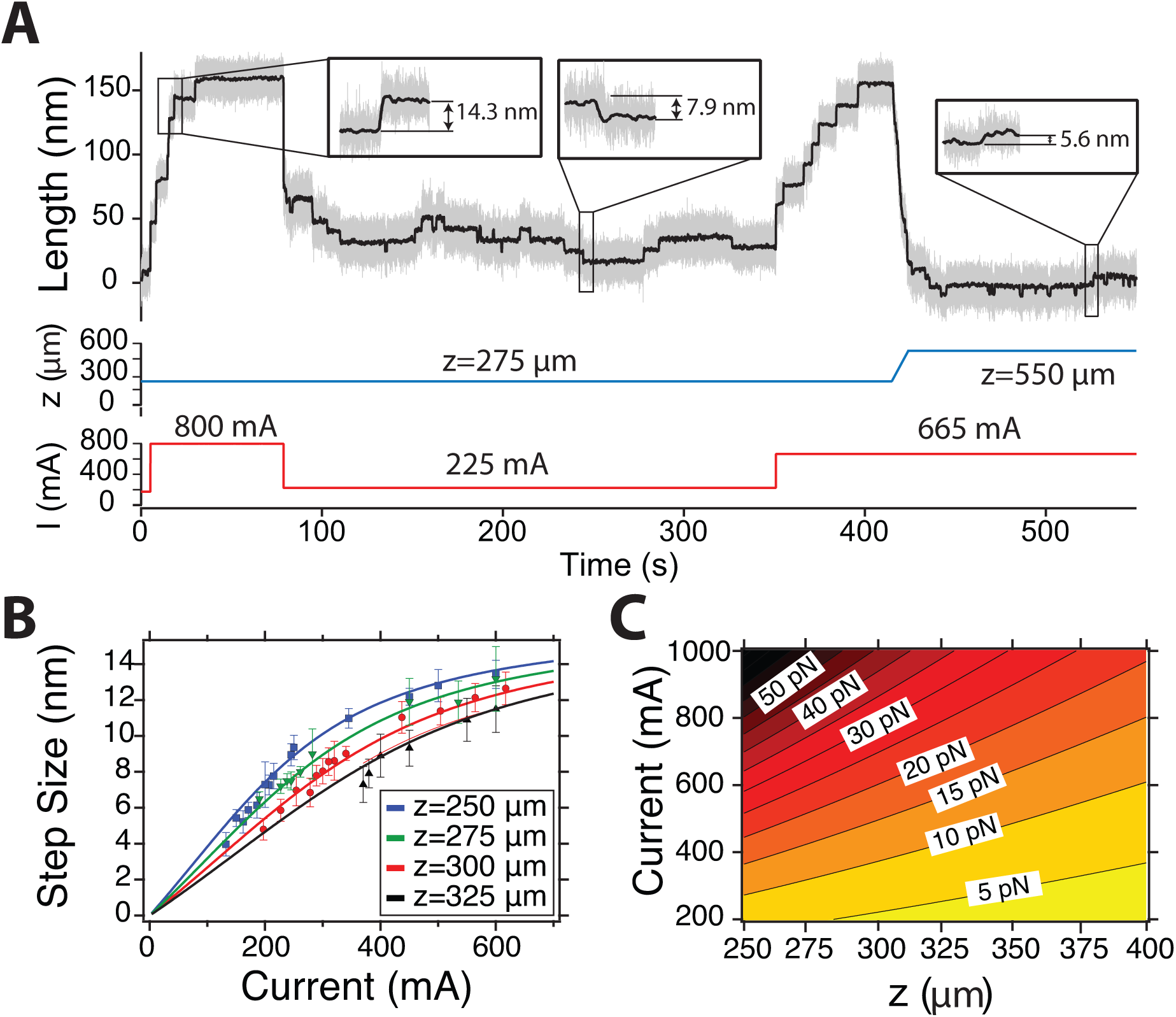
Calibration of the magnetic tape head tweezers. (A) Dynamics of protein L at different values of the current and vertical distance *z*. Data acquired at 1.4 kHz (grey trace), while the black curve is a filtered trace with a Savitzky-Golay algorithm (box size, 101). (B) Global fit of the step sizes to the FJC model to determine the current calibration law. Data collected over ¿40 molecules, and ¿300 events at each distance *z*. (C) Contour plot of the pulling force applied by the magnetic head to the superparamagnetic beads as a function of the vertical distance *z* and supplied current *I*.

Following this strategy, we measure folding/unfolding steps of protein L at distances between 250 *µ*m and 325 *µ*m, and currents between 200 and 1000 mA, since the magnetic head saturates at *I* = 1200 mA. Figure 2B shows the step sizes measured at four different distances, and plot as a function of *I*. Experimental data is fit globally to the FJC with *A* and *B* as only free parameters using the Levenberg-Marquardt algorithm, yielding *A* = 0.386 ± 0.131 pN/mA^2^ and *B* = 9.462 ± 1.564 pN/mA. The fit was weighted with the standard deviation of the distribution of step sizes. Thus, we establish our current law to be 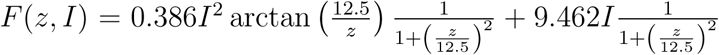, where *F* is given in pN, and *z* in *µ*m. This reveals that the dependence of the pulling force with *z* changes over a length scale of *g* = 25 *µ*m, illustrating the need of positioning the magnetic head with *µ*m precision, as any error in establishing *z* will result in a poor determination of *F*. We demonstrate the goodness of our calibration by reproducing the folding/unfolding properties of protein L (SI Appendix, Fig. S9), which match those previously measured [10, 24]. As we demonstrated previously [11], the bead-to-bead variation in the M-270 is very small (just 1.4% of dispersion in size, see SI Appendix). This allows to generate highly reproducible pulling forces within a 5% error, given the same magnetic gradient.

Figure 2C shows a contour plot of the force versus *z* and *I*, where colors scale from yellow to dark red as the force increases. The force depends in a strongly non-linear way on *z* and *I*, providing a convenient versatility in the control of the force, depending on the range we wish to work on. With our fluid chamber design (see SI Appendix), we can work at a minimum distances of *~* 250 *µ*m, which generates a maximum force of 50 pN. This range can be extended with alternative designs to produce thinner chambers.

### Verification of the calibration law using talin fine force sensitivity

We validate our description of the pulling force by confirming the adequacy of the theoretical framework we employed to predict the magnetic force on a superparamagnetic bead. In order to do so, we use the R3 IVVI mutant talin domain, given its extreme sensitivity to the force [25]. Figure 3A shows a trajectory of this talin monomer at three different forces (see Materials and Methods, and SI Appendix, Fig. S10A for the molecular construct), which highlights its aforementioned force sensitivity. Over just 0.8 pN, it transitions from an 80% probability of occupying the folded state (at 8.2 pN), to 20% at 9 pN, while at 8.7 pN, both states are equally populated. Figure 3B shows the folding probability calculated from length histograms of the time series, illustrating how subtle changes in the value of the force result in considerable shifts of the folding probability. This dependency has been independently measured with our standard magnetic tweezers setup (see SI Appendix, Fig. S10B).

**FIG. 3.**
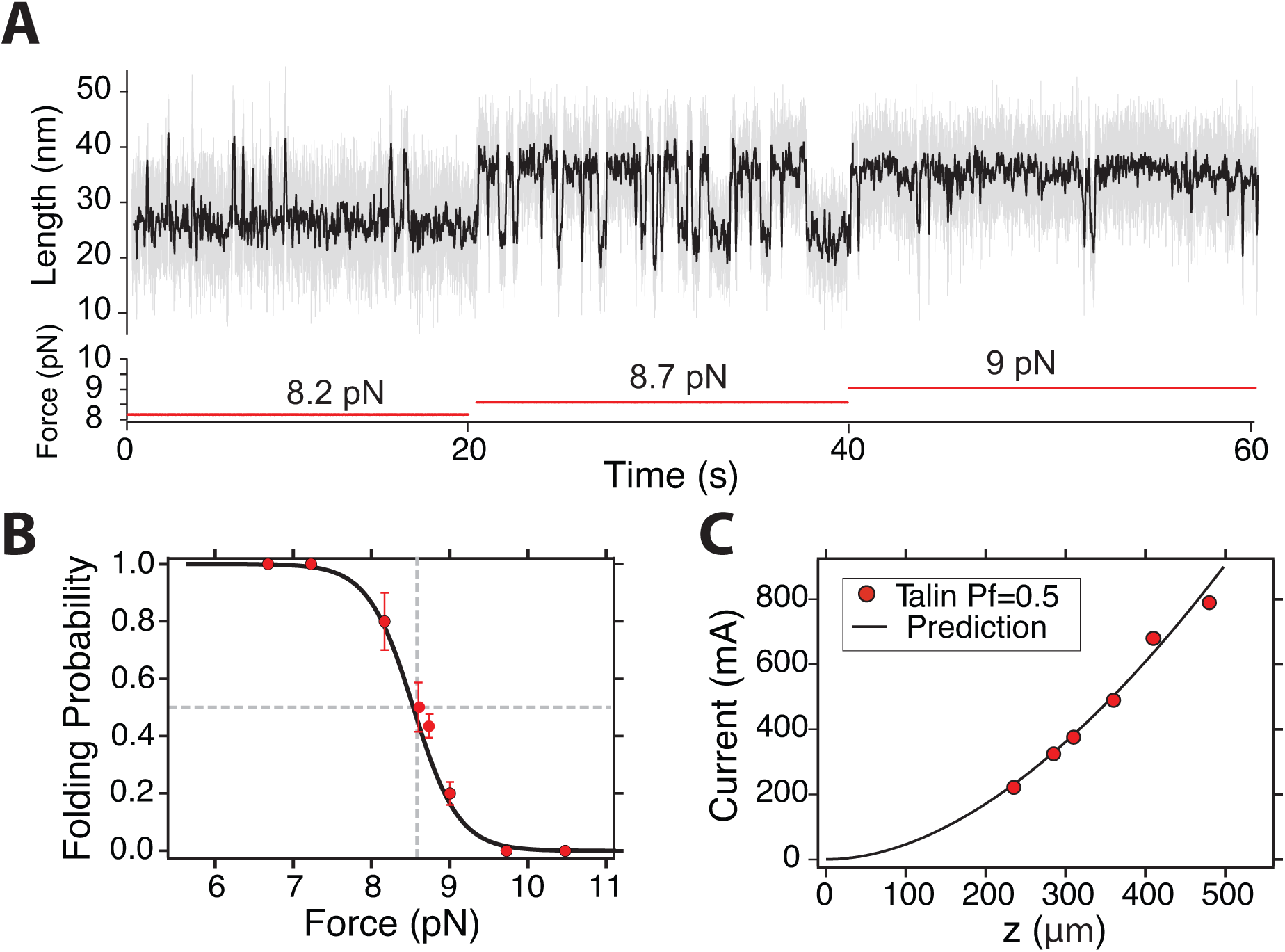
Talin as a molecular magnetometer. (A) Typical talin domain R3 trajectory at three different forces (8.2, 8.7, and 9 pN), sampled at 1.3 kHz. (B) Folding probability of talin measured with the magnetic tape head tweezers. Data measured with 4 molecules. (C) Electric current necessary to apply 8.7 pN at different values of *z* (manifested as *P_f_* = 0.5), together with the analytic prediction *I*(*z*).

In Fig. 3C, we plot the value of the electric current necessary to generate 8.7 pN (50% folding probability on talin) at different values of *z*. We can describe this dependence analytically by inverting Eq. 1, and obtaining *I*(*z*), which we plot as a solid black line in Fig. 3C (see SI Appendix, Eq. S23). Our analytic expression accurately follows the experimental data over a range *>* 250 *µ*m, providing a direct demonstration of the adequacy of Karlqvist description for the magnetic field created by the tape head, and the approximation for the force acting on a superparamagnetic bead [21].

### Direct detection of ephemeral states in the folding pathway of protein L

Our instrument allows to carry out very brief quench pulses to well-defined forces, and thus to directly sample the pathway of a folding protein with an unprecedented resolution. Here, we use it to explore the folding mechanism of protein L at a timescale of milliseconds. To this aim, we design the force protocol depicted in Fig. 4A. After the fingerprint pulse (i), we decrease the force to 10 pN to set a reference extension of the unfolded protein (ii), and subsequently to a folding force of 1 pN during varying times ∆*t*, ranging from 0.01 s to 10 s (iii). Next, we increase the force to 10 pN again (iv), which is the lowest force at which protein L does not fold, allowing us to observe any structure formed during the quench time with the slowest possible unfolding kinetics. This protocol is only possible because the magnetic head changes the field in *∼* 100 microseconds, so we can carry out controlled pulses of only few milliseconds long.

**FIG. 4.**
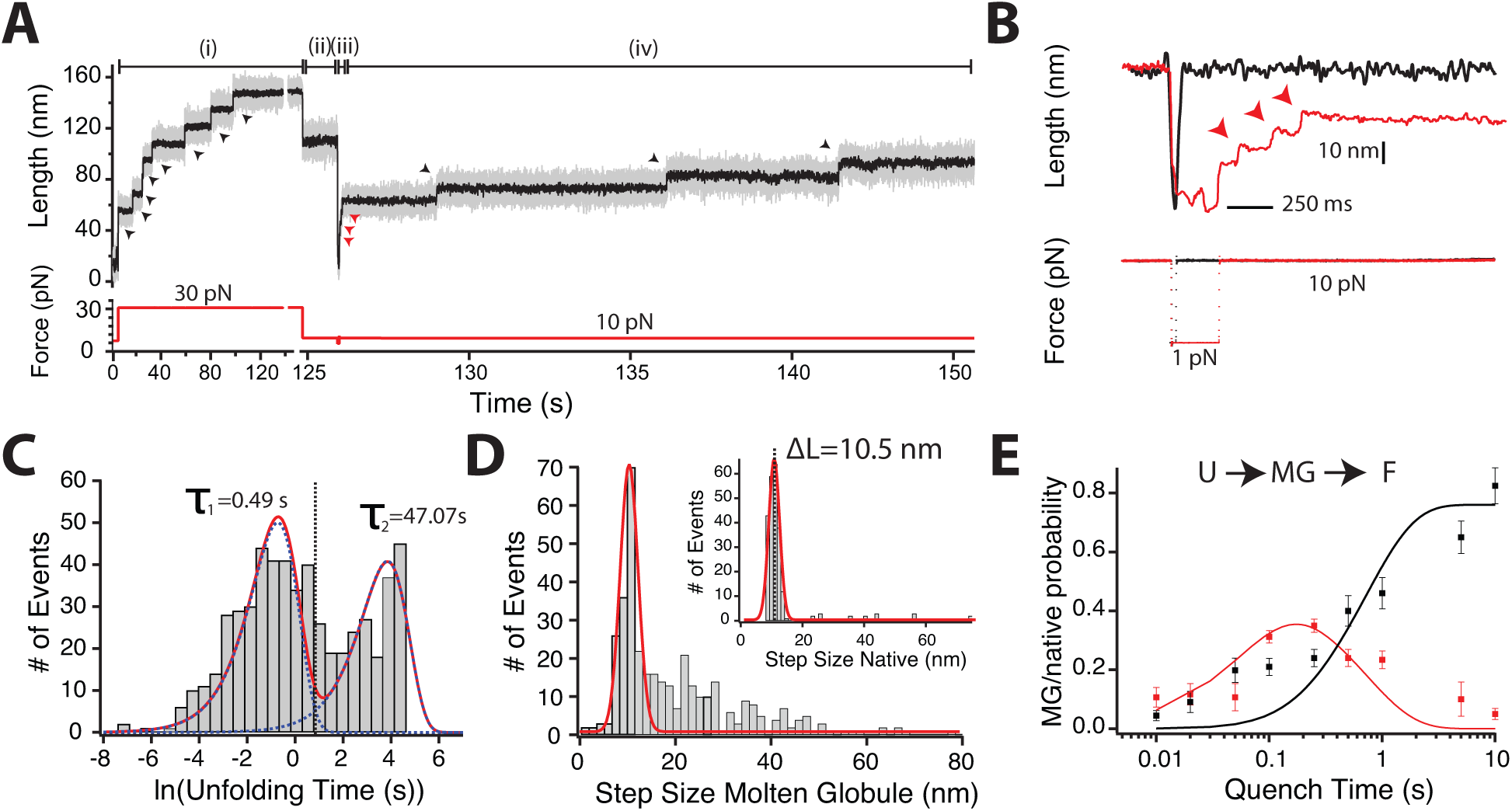
Molten globule state formation at a millisecond timescale. (A) Pulse protocol for characterizing the folding mechanism of protein L, sampled at 1.3 kHz. (B) Detail of the quench pulse, highlighting the formation of molten globule states over few milliseconds. (C) Square root histogram of the unfolding kinetics of all observed unfolding events at 10 pN, demonstrating the existence of two different populations. (D) Histogram of the unfolding step sizes of the weak domains (unfolding within less than 2.05 s), and the native states (inset). Histogram built from the aggregated data at every quench duration. (E) Probability of the molten globule (red) and native states (black) as a function of the quench time ∆*t*, globally fit to a first-order kinetics model with one intermediate. Data collected with 14 molecules, and a total of 546 events.

Figure 4B highlights the quench pulse for ∆*t* = 10 ms (black) and ∆*t* = 250 ms (red). The lower panel shows a direct measurement of the pulling force, obtained by sampling the electric current across the head at 100 kHz. The stepwise change from 10 to 1 pN is done with great precision and in less than 100 *µ*s (see SI Appendix, Fig. S7). This time constant depends on the power at which we supply current to the head, 60 W with the current implementation. When ∆*t* = 10 ms (black trace), the protein does not acquire any stable conformation, and the extension reaches the same level at 10 pN. However, when ∆*t* = 250 ms, we observe some ephemeral states that unfold in a sub-second timescale (red arrows), while those domains which acquired the native state unfold over several seconds (black arrows, Fig. 4A).

To characterize this folding mechanism in detail, we measure the unfolding kinetics of all unfolding events, and plot the histogram with logarithmic binning (square-root histogram, Fig. 4C). This representation allows for a systematic identification of different characteristic timescales, since an exponential distribution—signature of first-order kinetics—is written as *g*(*t*) = exp [*x − x*_0_ *−* exp(*x − x*_0_)] [26] (being *x* = ln *t*), where the average unfolding time *x*_0_ = ln *τ* is easily recognizable as a peak. Our distribution of unfolding events exhibits an evident bimodal shape (Fig. 4C), revealing the presence of two different kinetics. Fitting to a double exponential (red solid line), we identify a first set of ephemeral domains that unfold on average at *τ*_MG_ = 0.49 ± 0.01 s, and a more stable one where *τ*_N_ = 47.07 ± 14.511 s. This allows us to discriminate systematically between two kinds of events, setting a threshold time at 2.01 seconds (black dashed line). We measure separately the step sizes of the weak (Fig. 4D) and stable domains (Fig. 4D, inset). While the weak domains exhibit a broad step size distribution, the stable ones show a gaussian distribution centered at 10.5 nm. However, nearly 60% of the weak events fit onto an analogous step size distribution about 10.5 nm (solid red line), which suggests that they correspond to an immature conformation that eventually evolves to the stable native state (molten globule-like).

Our results suggest that protein L folds through a sequential reaction from the unfolded state (U) to a molten globule-like state (MG) that matures eventually to the native conformation (N). Figure 4E shows the probability of appearance of the molten globule and native conformations as a function of ∆*t*. The molten globule is manifested optimally in a timescale of *∼*100 ms, decaying quickly when we quench during seconds. Oppositely, the native domains show a sigmoid-like behavior, which saturates after few seconds. This dependence can be described analytically by a kinetic model where the molten globule is an intermediate of the native conformation (see SI Appendix for derivation). Solid lines show the global fits of the molten globule/native probability as a function of ∆*t* which yield the microscopic rates of formation of the molten globule (*r*_MG_ = 10.97 ± 1.42 s^*−*1^) and the native state (*r*_N_ = 1.28 ± 0.24 s^*−*1^). The fraction of native states does not saturate to 1 even when ∆*t* = 10 s, but rather to *∼* 0.8, and similarly the fraction of unfolded states plateaus at *∼*0.2 (see SI Appendix). This is due to the residual formation of stable non-native conformations with large step sizes (see inset in Fig. 4D), that can be related to domain swap-like structures [27].

Unlike previous observations of molten globule-like states in protein folding under force [28, 29], our instrumental design maintains a fine control of the pulling force, and allows measuring its evolution at different folding forces. With the protocol described in Fig. 4A, we keep the quench time constant at ∆*t*=250 ms and compare the formation of the molten globule at forces of 0.5, 1, 2, and 4 pN. Every of these forces produce observable molten globule-like states, which unfold within the first 2 seconds of the 10 pN probe pulse. However, the distribution of their step sizes is surprisingly dependent on the quench force. Figure 5 shows histograms of step sizes at 0.5, 1, 2, and 4 pN, normalized to facilitate their comparison. At forces equal or below 2 pN, we find a broad distribution, with a large fraction of steps that exceed a single native protein L. However, when we quench at a higher force of 4 pN, the distribution gets significantly narrower, and we observe very few large steps. Fitting gaussian distributions around 10.5 nm (red solid lines), we obtain respectively, that at, 0.5, 1, and 2 pN, 45%, 55% and 48% of the steps are short, while at 4 pN, this fraction increases to 75%. This observation suggests that mechanical forces may assist folding by facilitating the formation of molten globule-like states that lead to the native state, in contrast to the longer collapsed structures that hamper the folding reaction. Indeed, our data indicates that the probability of reaching the native state after a brief 250 ms long quench increases with the force; while at 0.5, 1 and 2 pN only 25% of the domains reached the native conformation, a higher force of 4 pN allowed 50% of the domains to fold (see Table S3).

**FIG. 5.**
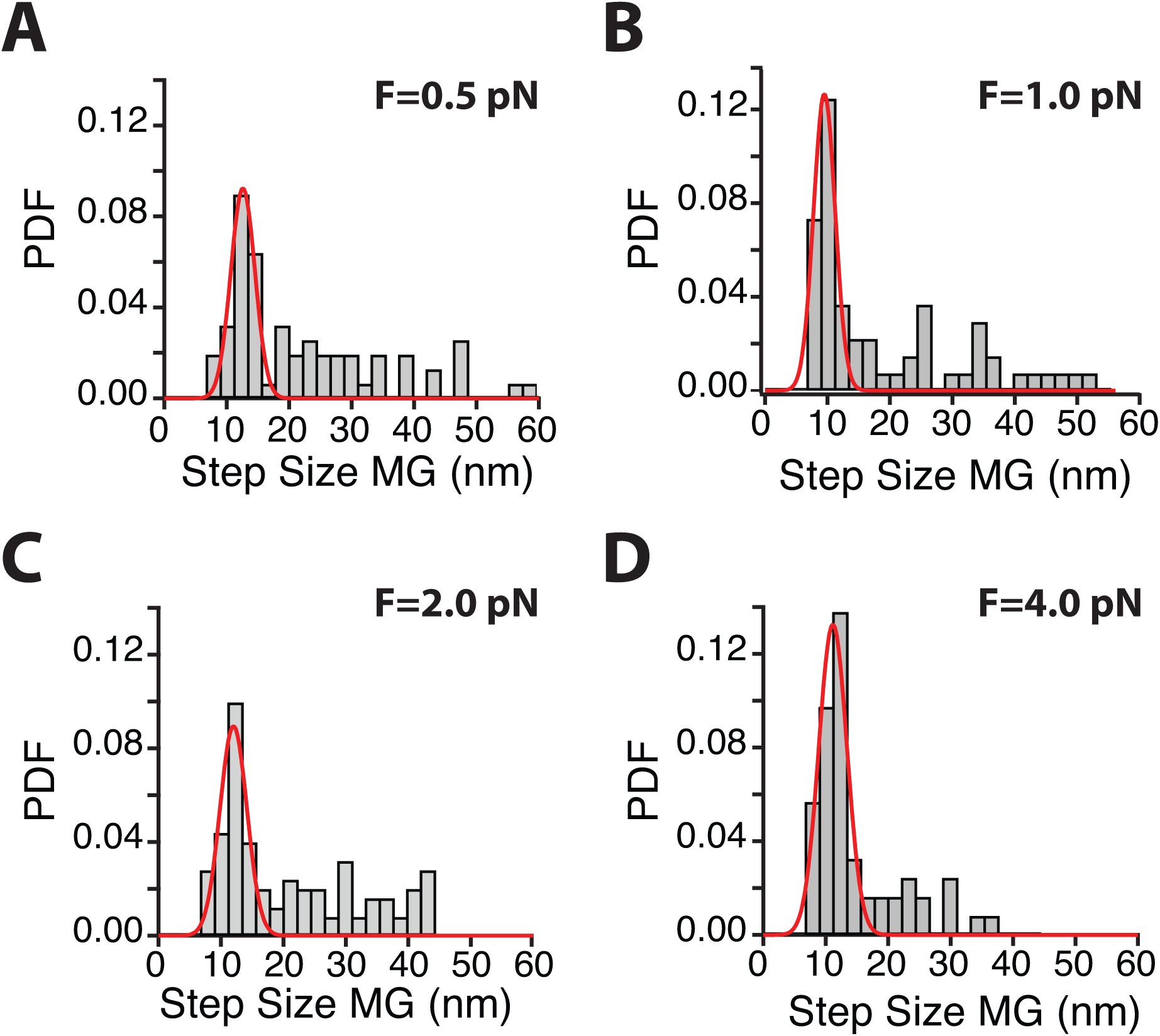
Probability distribution function (PDF) of the step sizes of the ephemeral states measured at quenching forces of 0.5 pN (A), 1.0 pN (B), 2.0 pN (C) and 4.0 pN (D). Data at 1 pN corresponds to the 250 ms long quenches from Fig. 4D. Red line is a gaussian fit to the molten globule-like states, peaked at 10.5 nm. As we increase the quench force the distribution narrows, and the probability of finding large steps decreases as the force increases. At 0.5, 1 and 2 pN, 45%, 55% and 48% of the ephemeral states are distributed around 10.5 nm, while this fraction increases to 75% at 4 pN. The distributions were measured with 71 (0.5 pN), 62 (1 pN), 113 (2 pN), 61 (1 pN) and 56 (4 pN) events, and in a total of 8 molecules.

## DISCUSSION

It is now well understood that upon quenching the force on a mechanically unfolded/extended protein, the polypeptide rapidly collapses, greatly increasing its entropy. For ubiquitin polyproteins studied with force-clamp AFM, this collapse was fast, taking place in the submillisecond time scale [28, 30, 31] The collapsed polypeptide was then observed to progress over a time scale of *∼*1 second to its native mechanically stable conformation [28]. This progression from collapsed-to-native proceeded through an ensemble of mechanically weak states that were mechanically labile and showed a wide distribution of step sizes. Akin to those transient folding states observed in bulk [32–34], these mechanically weak states were proposed to be molten-globule like states. The instrumental limitations of the force-clamp AFM were evident in those studies. The time response of the force-clamp apparatus was ¿3-5 ms and the quench force could not be ascertained with precision in the pN range [28, 30]. These limitations prevented an accurate study of the weak intermediate states, beyond noting their existence. Similarly, Cecconi and colleagues also observed the existence of a mechanically labile on-pathway molten globule-like state in RNase H [29] that, at a force of a few pN, matured slowly to its mechanically stable state. Again, the limitations of the optical trapping method were evident in these experiments where rapid force changes could not be implemented, thus the time/force evolution of molten globule states could not be studied.

The magnetic tweezers design that we present here overcomes these issues by demonstrating an exquisite control of the pulling force in the pN range combined with a bandwidth better than 10 kHz. We have applied our novel force spectroscopy technique to study protein L, a classical two state folder that is not known to have a molten globule state [35]. Protein L has been shown to be a mechanically weak protein that folds without showing obvious mechanical intermediates [10, 13]. Here, we discover that protein L folds through very short lived and mechanically weak intermediates (Fig. 4), similar to those observed earlier for ubiquitin [28], but on a much shorter time scale. One noteworthy result was the observation that the distribution of step sizes of the mechanically weak molten globule states varied significantly with the quench force. Histograms of unfolding step sizes of the molten globule states at 10 pN showed a narrowing distribution approaching that of the native conformation, as the quench force was increased up to 4 pN (Fig. 5). Our results suggest that quenching to lower forces (e.g. 0.5 pN; Fig. 5A) drives a collapsing polypeptide towards more disordered states. It is striking that quenching to higher forces results into a much narrower distribution of molten globule states (Fig. 4), and a higher probability of maturation to the native conformation. These results suggest that there is an optimal force for the formation of the molten globule states that lead to the native state. Highly disordered collapsed states, such as those that form at the lowest forces, may have such a high entropy that makes it more difficult to sort out the correct positioning of the polypeptide to assemble the native state. The instrument that we demonstrate here now opens a path for detailed studies of such weak ephemeral states in protein folding.

In summary, we have implemented a magnetic tape head in a novel force spectrometer, which enables to change the force with a bandwidth above 10 kHz, while maintaining high accuracy and stability. The force generated by the head can be described analytically with great accuracy, and over a large range of forces. The unique capability of changing the force in few microseconds has allowed us to directly observe for the first time the molten globule state in protein L, which forms in few milliseconds and would be otherwise hidden in the dead time introduced by force spectrometers that change the force mechanically. Furthermore, magnetic tape head tweezers opens the possibility of novel force protocols, more complex than classic square pulses, or ramps. Force spectroscopy has been traditionally used to investigate how do proteins respond to pulling forces of different magnitude. However, it remains unexplored how fast do proteins respond to forces that change in time, or to signals with different spectral densities. Mechanical forces in physiology are likely to be embedded in mechanical noise [36, 37], and to contain oscillatory components [38–40], but the effect of these signals on protein folding remains unknown. We envision that magnetic tape head tweezers will be able to address these problems by recreating mechanical perturbations which mimic those encountered in physiology, and thus to open a range of unexplored biological problems.

## MATERIALS AND METHODS

### Magnetic tape head tweezers setup

All experiments were carried out in a custom-made setup built on top of an inverted microscope, with a piezo-mounted objective (P-725; Physik Instrumente). Images were acquired using a CMOS Ximea MQ013MG-ON camera. The mechanical force was applied on superparamagnetic beads Dynabeads M-270 beads through a magnetic tape head (BRUSH 902836) mounted on an articulating arm and positioned with micrometer resolution. The data acquisition and control of the piezo actuator was done using a multifunction DAQ card (NI USB-6289, National Instruments). Image processing was done by a custom-written software written in C++/Qt, fully available in the lab’s website (http://fernandezlab.biology.columbia.edu/). For further details see SI Appendix.

## Single molecule measurements

All experiments were carried out in custom-made fluid chambers (see SI Appendix), functionalized as described before [6]. All polyprotein constructs were flanked by an N-terminal HaloTag enzyme (Promega) and a C-terminal AviTag for biotinylation. For detailed expression and purification methods see SI Appendix. All experiments were done in HEPES buffer (Hepes 10 mM pH 7.2, NaCl 150 mM, EDTA 1 nM) containing 10 mM ascorbic acid (pH 7.3) to avoid oxidative damage, using a protein concentration of *∼* 50 nM. Data was acquired at a rate between 1.2 and 1.7 kHz, and subsequently filtered using a Savitzky-Golay algorithm with a 101 points box, to facilitate visualization and analysis. Step sizes were measured using length histograms of the filtered trace. Molten globule experiments were analyzed using the raw trace, due to their fast timescale. Further details on experimental analysis are described in SI Appendix.

## ACKNOWLEDGMENTS

This work was supported by NIH Grants GM116122 and HL061228. R. T-R. acknowledges Fundacion Ramon Areces for financial support. E.C.E. was supported by NIH F30HL129662. We thank Dr. Igor Barsukov from University of Liverpool for sharing the talin plasmid with us. We would like to thank all the members of Fernandez lab for the valuable discussions and comments on the manuscript.

