## Supplementary material for "Ephemeral states in protein folding under force captured with a novel magnetic tweezers design": SI

### DERIVATION OF THE CURRENT LAW

The shape of the magnetic field created by a tape head can be obtained by solving the Maxwell equations with the appropriate geometry and boundary conditions. This problem is not trivial and has been tackled through different strategies, both analytical and numerical. Karlqvist approximation is the most widely used expression for the field generated by a magnetic tape head, and perhaps the simplest one [1]. Assuming a two dimensional problem, and that the gap surface potential is linear, the field can be written as

$$B_{\mathrm{x}} = \frac{B_{\mathrm{g}}}{\pi}\left[\arctan\left(\frac{g/2+x}{z}\right) + \arctan\left(\frac{g/2-x}{z}\right)\right], \tag{1}$$

$$B_{\mathrm{z}} = -\frac{B_{\mathrm{g}}}{2\pi}\ln\frac{(g/2+x)^2+z^2}{(g/2-x)^2+z^2}, \tag{2}$$

where $B$ is the magnetic field (T), $x$ is the lateral coordinate and $z$ the vertical one, $g$ is the width of the gap, and $B_{\mathrm{g}}$ the deep gap field, this is, the magnetic field *inside the gap*. For a generic magnetic head [2]:

$$B_{\mathrm{g}} = \mu_0 \frac{NI}{g}\eta, \tag{3}$$

where $N$ is the number of wire turns around the head core, $I$ the intensity of the electric current (A), $\mu_0 = 4\pi \times 10^{-7}$ Tm/A the permeability of the vacuum, and $\eta$ the field efficiency. The parameter $\eta$ is defined as the fraction of potential that reaches the gap and the fraction of signal flux that reaches the coil. For realistic heads, $\eta$ is in general very hard to calculate since it must account for the detailed geometry of the head [2]. For an idealized head, one can write:

$$\eta = \frac{g/A_{\mathrm{g}}}{g/A_{\mathrm{g}} + l_c/(\mu_{\mathrm{r}} A_{\mathrm{c}})}, \tag{4}$$

where $A_{\mathrm{g}}$ the sectional area of the gap, $\mu_{\mathrm{r}}$ the relative permeability of the medium and $A_{\mathrm{c}}$ the cross sectional area of the core [2]

Karlqvist approximation assumes a linear relationship between the magnetic filed and the electric current, and thus it is only applicable in the regime. If we assume a simplistic description of a magnetic head where the field increases linearly until it saturates, we could write $B_{\mathrm{g}} = B_{\mathrm{gs}} I/I_{\mathrm{s}}$, being $B_{\mathrm{gs}}$ the gap field at the saturation current $I_{\mathrm{s}}$. In our case, the manufacturer specifies a saturation current $I_{\mathrm{s}} = 1.2$ A, and a maximum gap field $B_{\mathrm{gs}} = 0.5$ T. However, this description of the field is a rough approximation, and the complete response of the head is unknown.

Figure 1 shows the $x$ (solid) and $z$ (dotted) components of the field with $B_{\mathrm{g}} = 500$ mT for three different working distances. Since $g = 25$ $\mu$m, the gap spans from -12.5 $\mu$m to 12.5 $\mu$m, and in that region $B_{\mathrm{x}}$ is practically constant, and $B_{\mathrm{z}}$ negligible, meaning that working under the gap would result in a single pulling force component in $z$.

The force acting on a bead can be modeled as:

$$\vec{F} = (\vec{m} \cdot \nabla) \vec{B}, \tag{5}$$

being $\vec{m}$ the magnetic moment of the bead (Am$^2$), and $\vec{B}$ the applied field. In the literature it is usual to find $\vec{F} = (V\Delta\chi/\mu_0)\left(\vec{B} \cdot \nabla\right)\vec{B}$ without derivation [3–7], where $V$ is the volume of the bead and $\Delta\chi$ the difference in magnetic susceptibilities between the particle and the medium (dimensionless). This expression assumes that the magnetization is linear with the applied field $m \propto B$, which is true at weak fields. If one assumes that the magnetization of the field follows a common Langevin-like dependence, $m = m_{\mathrm{s}} (\coth B/B_{\mathrm{s}} - B_{\mathrm{s}}/B)$, we can assume this linear dependence for $B < B_{\mathrm{s}}$.

Additionally, we can assume that the beads have a non-zero magnetization $m_0$ at zero field, even though our beads are superparamagnetic. Although bulk measurements typically demonstrate $m_0 = 0$ at zero field, the behavior of the individual beads can be different, and thus it is appropriate to include this term as a free parameter [8]. We will further discuss this issue in next section. The magnetization can be written as:

$$\vec{M} = \vec{M}_0 + \frac{\chi_{\mathrm{b}}}{\rho}\frac{\vec{B}}{\mu_0}. \tag{6}$$

where here $\vec{M}$ is the average mass magnetization of the beads (Am$^2$/kg), $\vec{M}_0$ the initial magnetization, $\chi_{\mathrm{b}}$ the initial susceptibility of the bead (dimensionless) and $\rho$ its density. Thus, the force acting on a superparamagnetic bead at low fields is:

$$\vec{F}_{\mathrm{m}} = \rho V \left(\vec{M}_0 \cdot \nabla\right) \vec{B} + \frac{V\chi_{\mathrm{b}}}{\mu_0}\left(\vec{B} \cdot \nabla\right)\vec{B}, \tag{7}$$

where $V$ is the volume of the bead. The limits of this approximation are discussed in Section II. Furthermore, we develop an expression based on a Langevin description of the magnetization in Section III and explore its applicability.

Now, considering a two-dimensional problem, and assuming Karlqvist approximation for the magnetic field, the magnetic force takes the explicit form:

$$\vec{F}_{\mathrm{m}} = \frac{V\chi_{\mathrm{b}}}{\mu_0}\begin{pmatrix} B_{\mathrm{x}}\frac{\partial B_{\mathrm{x}}}{\partial x} + B_{\mathrm{z}}\frac{\partial B_{\mathrm{x}}}{\partial z} \\ B_{\mathrm{x}}\frac{\partial B_{\mathrm{z}}}{\partial x} + B_{\mathrm{z}}\frac{\partial B_{\mathrm{z}}}{\partial z} \end{pmatrix} + \rho V \begin{pmatrix} M_{0\mathrm{x}}\frac{\partial B_{\mathrm{x}}}{\partial x} + M_{0\mathrm{z}}\frac{\partial B_{\mathrm{x}}}{\partial z} \\ M_{0\mathrm{x}}\frac{\partial B_{\mathrm{z}}}{\partial x} + M_{0\mathrm{z}}\frac{\partial B_{\mathrm{z}}}{\partial z} \end{pmatrix}. \tag{8}$$

Since we work under the gap region, $x \approx 0$, the x component of the force vanishes. Also, because the bead can rotate freely, we can assume that the $z$ component of the initial magnetization is zero, and thus $\vec{M}_0 = (M_{0\mathrm{x}}, 0)$. Under these condition the $x$ component of the force vanishes, and we can write:

$$F_{\mathrm{z}} = \frac{V\chi_{\mathrm{b}}}{\mu_0} B_{\mathrm{x}} \frac{\partial}{\partial x} B_{\mathrm{z}} + \rho V M_{0\mathrm{x}} \frac{\partial}{\partial x} B_{\mathrm{z}}, \tag{9}$$

where:

$$B_{\mathrm{x}} = 2\frac{B_{\mathrm{g}}}{\pi} \arctan\left(\frac{g/2}{z}\right),$$

and

$$\frac{\partial}{\partial x} B_{\mathrm{z}} = \frac{B_{\mathrm{g}}}{\pi}\frac{4}{g}\frac{1}{1+\left(\frac{z}{g/2}\right)^2}.$$

Thus, the force acting on the superparamagnetic bead (being a pulling force in the $z$ direction):

$$F(z, I) = AI^2 \arctan\left(\frac{g/2}{z}\right)\frac{1}{1+\left(\frac{z}{g/2}\right)^2} + BI\frac{1}{1+\left(\frac{z}{g/2}\right)^2}, \tag{10}$$

where the constants $A$ and $B$:

$$A = 8V\chi_b\mu_0\frac{N^2}{g^3}\frac{\eta^2}{\pi^2} \tag{11}$$

$$B = 4\rho V\mu_0\frac{N}{g^2}\frac{\eta}{\pi}M_{0\mathrm{x}} \tag{12}$$

We leave both $A$ and $B$ as free parameters, specific for the calibration given the magnetic head (Brush Industries, 902836) and beads (Dynabeads M-270$^{\mathrm{TM}}$) we use.

Next, since the dependence of the step sizes with the pulling force $F(z, I)$ can be described by the freely jointed chain model:

$$\Delta L = \Delta L_{\mathrm{c}} \left[ \coth\left( \frac{F(z,I) l_{\mathrm{K}}}{kT} \right) - \frac{kT}{F(z,I) l_{\mathrm{K}}} \right], \tag{13}$$

we can determine $A$ and $B$ by measuring $\Delta L$ as a function of $z$ and $I$ and fitting globally to Eq. 13, where $\Delta L_{\mathrm{c}} = 16.3$ nm and $l_{\mathrm{K}} = 1.1$ nm, for protein L [9].

### MAGNETIC CHARACTERISTICS OF THE BEADS AND LIMITS OF APPLICABILITY OF THE CURRENT LAW

We discuss here the magnetic properties of the Dynabeads™M-270 superparamagnetic beads, employed throughout this work. This allows us to explore the range of validity of our current law. We summarize the magnetic properties of these beads in Table I, as described in [10].

Our current law was developed assuming a linear dependence between the magnetization $\vec{M}$ and the applied field $\vec{B}$, including a term for the initial magnetization of the bead (Eq. 6). This approximation is valid at low fields. If we assume a typical magnetization curve, which follows a Langevin-like dependence:

$$M(B) = M_s \left[ \coth\left( \frac{B}{B_s} \right) - \frac{B_s}{B} \right], \tag{14}$$

the magnetization is linear with the field for $B < B_{\mathrm{s}}$, and it saturates to $M = M_{\mathrm{s}}$. For the M-270 beads, the magnetization-field relationship was measured in [10], which we reproduce in Fig. 2, and fit to Eq. 14. We obtain $M_{\mathrm{s}} = 10.3$ Am$^2$/kg and $B_{\mathrm{s}} = 19$ mT. Thus, we can consider that our approximation is valid if the field on the bead is below 19 mT, approximately. This is in agreement with that discussed in [8], where they use the same approximation on different superparamagnetic beads for fields below 3 mT, at which the magnetization is approximately $0.2M_{\mathrm{s}}$.

We can now calculate the field on the beads, by assuming $B_{\mathrm{g}} = B_{\mathrm{gs}} I/I_{\mathrm{s}}$, which is a rough approximation of the field on the beads, but provides us an estimation of the field on the beads in the z-I regime we work on. We can plot $I(z)$ constant field lines as:

$$I = \frac{\pi B I_{\mathrm{s}}}{2B_{\mathrm{gs}}} \frac{1}{\arctan\left(\frac{g/2}{z}\right)} \approx \frac{\pi B I_{\mathrm{s}}}{2B_{\mathrm{gs}}} \frac{z}{g/2}, \tag{15}$$

since $\arctan(x) \approx x$ for small $x$, and $(g/2)/z \ll 1$ in our experimental applications, since g=25 $\mu$m

Figure 3 shows the $I$ versus $z$ curve for different values of the magnetic field, with the current-distance working region highlighted in green. In the region where we have calibrated our setup (250 $\mu$m < z < 400 $\mu$m; and $0 < I < 1$ A), the field on the bead is < 15 mT. Thus, the linear approximation is valid.

Moreover, Eq. 10 includes a term to model the initial magnetization of the bead $M_0$. Superparamagnetic beads should ideally have $M_0 = 0$. This is shown in the hysteresis $M(B)$ curves, as that measured in [10] for the M-270 (from which the data shown in Fig. 2 was taken). However, this measurements are typically done by preparing a sample in the form of dry powder, with millions of beads, which have random orientations. Therefore, the initial magnetization of the sample is zero, which does not imply that $M_0$ for the individual beads is zero. This issue is discussed in Ref. [8], where they argue that, at weak fields, the initial magnetization can take values comparable to the induced one. Indeed, they demonstrate that a non-zero $M_0$ is necessary to correctly model the force acting on individual superparamagnetic beads, and that that initial magnetization can take values as large as $0.07M_s$. This justifies that our force equation Eq. 10 depends on two terms, one given by the induced magnetization, and a second one given by the initial magnetization.

We can estimate how important are relatively these two terms. When both contributions are equal, the field on the beads is $B = 6.6$ mT (this is easily estimated by calculating the I(z) relation where both contributions are equal, and calculating the field through Eq. 15). At this field, the induced magnetization on the beads is $\sim 0.11M_{\mathrm{s}}$, meaning that our model suggests $M_0 \sim 0.11M_{\mathrm{s}}$, which is in the range demonstrated in [8].

##### Limitations of the model and consistency check

Here, we gather the assumptions of the model and its range of applicability, and check the consistency of the values obtained from the calibration of our instrument, given the physical characteristics of our problem.

Equation 10 is assumes that the magnetic field created by the tape head can be modeled by Karlqvist approximation (Eqs. 1 and 2), and that the magnetization of the bead follows Eq. 6. Karlqvist approximation assumes a simplified description of a tape head, such as that it consists of two semi-infinite poles, which are made of a very soft magnetic material with infinite permeability, and that such poles are separated by a gap over which the field is constant, among others [1]. Additionally the field in the gap $B_{\mathrm{g}}$ increases linearly with the electric current $I$, which, for a realistic tape head, is true only for a certain range of currents. In our particular case, the saturation current is 1.2 A, but, as indicated by the manufacturer, the gap field is linear only up to 1 A. The validity of Eq 6 was already discussed in Section II. In this sense, our calibration is valid when using the tape head model from Brush industries (BRUSH 902836) and M-270 superparamagnetic beads from Dynabeads$^{\mathrm{TM}}$, and working with currents below 1 A, and at distances above 250 $\mu$m. In such case, the pulling force on the bead can be accurately described by the fitting parameters:

$$A = 0.386 \pm 0.131 \text{ pN/mA}^2$$
$$B = 9.462 \pm 1.564 \text{ pN/mA}.$$

Our calibration method using the step sizes of protein L is completely general, and new parameters $A$ and $B$ could be determined in case a different head or beads are employed, given that the conditions for using our model apply.

Equations 11 and 12 express $A$ and $B$ in terms of the physical parameters of the system. However, the geometrical parameters from the head $N$ and $\eta$ are unknown. We can assume, that roughly, the field increases linearly till $I_{\mathrm{s}} = 1.2$ A, where it saturates to $B_{\mathrm{gs}} = 0.5$ T, so that we could write

$$A \approx 8V\chi_{\mathrm{b}} \frac{1}{\pi^2 \mu_0 g} \frac{B_{\mathrm{gs}}^2}{I_{\mathrm{s}}^2},$$
$$B \approx 4\rho V M_{0\mathrm{x}} \mu_0 \frac{1}{\pi g} \frac{B_{\mathrm{gs}}}{I_{\mathrm{s}}}.$$

We can use the value we obtain for $A$ from our calibration to estimate the magnetic susceptibilities of the beads, which yields $\chi_{\mathrm{b}}^{\mathrm{model}} \approx 7.7$, which is approximately one order of magnitude higher than the actual one. Additionally, from the value of $B$ we can estimate the initial magnetization of the beads, which yields $M_{0\mathrm{x}}^{\mathrm{model}} \approx 26.8\ \mathrm{Am^2kg^{-1}}$, approximately a factor of 20 higher than what expected.

However, these values are obtained assuming a linear relation between $I$ and $B$ over the whole range of currents, and that $B_g = 0.5$ T at a current of 1.2 A, which is a rough approximation of the behavior of the head. A more realistic description of the field-current relation would likely decrease the estimation of $\chi_{\mathrm{b}}$ and $M_{0\mathrm{x}}$. Nevertheless, our model provides an unsatisfactory estimation of the parameters of the system, and the origin of such discrepancies, is hard to identify, given the range of approximations we have used. Regardless, our model provides a correct description of the pulling force as a function of the current and distance, which allows to accurately predict the pulling force over the current and distance regime we work on.

### DERIVATION OF THE CURRENT LAW FROM LANGEVIN DESCRIPTION OF THE MAGNETIZATION

Here, we develop an discuss an alternative expression for our current law, based on a Langevin description of the magnetization, where we assume that the magnetic moment of the bead:

$$m_{\mathrm{x}} = m_{\mathrm{s}} \left( \coth \frac{B_{\mathrm{x}}}{B_{\mathrm{s}}} - \frac{B_{\mathrm{s}}}{B_{\mathrm{x}}} \right). \tag{16}$$

Thus, the force acting on the bead would be described as:

$$\vec{F} = (\vec{m} \cdot \vec{\nabla}) \vec{B} = m_{\mathrm{x}} \begin{pmatrix} \frac{\partial}{\partial x} B_{\mathrm{x}} \\ \frac{\partial}{\partial x} B_{\mathrm{z}} \end{pmatrix}. \tag{17}$$

Given the Karlqvist description of the magnetic field, at $x = 0$, $\partial B_{\mathrm{x}}/\partial x$ vanishes, so we just have the $z$ component, and

$$\frac{\partial}{\partial x} B_{\mathrm{z}} = -\frac{4B_{\mathrm{g}}}{\pi g} \frac{1}{1 + z_{\mathrm{g}}^2}, \tag{18}$$

where $z_{\mathrm{g}} = 2z/g$. Thus, the force on the superparamagnetic bead:

$$F_{\mathrm{z}} = \frac{4B_{\mathrm{g}}}{g\pi} \frac{m_{\mathrm{x}}}{1 + z_{\mathrm{g}}^2}, \tag{19}$$

where we have dropped the $-$ sign, assuming implicitly that the force is a pulling force. Equation 19 is the force acting on a superparamagnetic bead assuming Karlqvist approximation for the magnetic field, and that the magnetization of the bead can be described by Langevin approximation.

If the force is to be expressed in pN, and $B_\mathrm{g}$ in T then:

$$F_\mathrm{z} = 10^{12} \frac{4B_\mathrm{g}}{\pi g} \frac{m_\mathrm{x}}{1+z_\mathrm{g}^2}, \tag{20}$$

where $[g] = \mathrm{m}$ and $[m_\mathrm{x}] = \mathrm{Am}^2$.

### CALIBRATION OF THE MAGNETIC TAPE HEAD TWEEZERS USING LANGEVIN APPROXIMATION FOR THE MAGNETIZATION OF THE BEAD

We calibrate our instrument with the step sizes of protein L at different distances and currents, using Eq. 13, where the force relation is given by Eq. 19. The parameters of this force law are $m_\mathrm{s}$, the saturation magnetization of the bead; $B_\mathrm{s}$, the field at which magnetization of the bead is $\sim 0.3m_\mathrm{s}$; $H_\mathrm{gs}$, the maximum gap field of the head; $I_\mathrm{s}$ the saturation current of the bead, and $g$ the gap width.

The parameters regarding the tape head are $B_\mathrm{gs} = 0.5$ T, $I_\mathrm{s} = 1.2$ A, and $g = 25\ \mu$m. The parameters of the beads, according to [10], are $B_\mathrm{s} = 0.019$ T, and $m_\mathrm{s} = 214.36 \times 10^{-15}$ Am$^2$.

We fit Eq. 13 to our step sizes, first just leaving $m_\mathrm{s}$ free, and then $m_\mathrm{s}$ and $B_\mathrm{gs}$ (Figure 4). The fits produced are very similar, and, attending to the residuals, are clearly skewed at large currents. The first fit yields of $m_\mathrm{s} = 6.65 \times 10^{-12}$ Am$^2$ which is about one order of magnitude larger than that suggested by the manufacturer. The second fit yields $m_\mathrm{s} = 4.48 \times 10^{-13}$ Am$^2$, closer to the actual one, and $B_\mathrm{gs} = 2$ T, which is overestimated by a factor of 4. In this sense, and like the model we employ, this description is not completely consistent with the known values of of the magnetic head and beads, as provided by the respective manufacturer.

However, the most important issue of the present model is that it does not provide an accurate description of the pulling forces, since both fits are biased at high currents. For instance, at a distance of 275 $\mu$m and a current of 600 mA, it predicts that the pulling force is 35 pN, when the measured step size is $\sim$ 13nm corresponding to a force of $\sim$ 20 pN.

Thus, considering that the ultimate objective of this model is to provide a calibration of our instrument, it is clear that this strategy overestimates the force at large values of the current, likely due to the intrinsic nonlinearity of the Langevin equation, compared to the linear approach, which is indeed valid in the current/distance regime we are working on.

### MAGNETIC TAPE HEAD SPECIFICATIONS AND IMPLEMENTATION DETAILS

We employ a commercially available magnetic tape head from Brush Industries (BRUSH 902836). The technical specifications are summarized in Table II.

The resistance of the head increases with the current since the head heats up. We measure the resistance of the head using a calibrated current source and measuring the voltage drop across the head (Figure 5, red points V(I), inset shows the resistance in Ω), and it ranges from $\sim 1.6\ \Omega$ at very low currents to $\sim 2.2\ \Omega$ at 2 A.We can model the dependence of the current as $R(I) = R_0 + kI^2$, thus $V_\mathrm{H} = R_0 + kI^3$. Fitting to this curve (solid red line) yields $R_0 = 1.65\ \Omega$ and $k = 0.13\ V/A^3$. In this sense, it is not practical to measure the voltage across the head to determine the current, given the variability of its resistance. Therefore, it is not practical to sample the current through the head directly.

To solve this problem, we introduce a precision resistance of 2 Ω connected in series with the head, and calculate the current flowing through the head by measuring the voltage drop across the measuring resistance. This measuring resistance is large enough to dissipate the heat and its value does not change with the current. We measure the voltage drop at different currents (Figure 5, blue), and it is linear with a slope of $R_\mathrm{M} = 2.0124\ \Omega$, with currents up to 2 A. Figure 6 shows a diagram of the current control circuit for the head.

### DIRECT MEASUREMENT OF THE CURRENT CHANGE

In order to demonstrate that our implementation of the instrument allows to change the force acting on the bead within 100 $\mu$s, we measure directly the electric current across the tape head during a current step, sampled at a rate of 100 kHz. Figure 7 shows this measurement for a stepped change in current between 345 mA and 102 mA. At a distance of 300 $\mu$m, this corresponds to a change in force between 9 pN and 2 pN. This illustrates that the step occurs within 100 $\mu$s, and the effect of the PID in the relaxation of the electric current is observed as a damped oscillation that equilibrates quickly. The electric current is directly transformed on a magnetic field gradient, and thus onto the pulling force on the bead.

### TEMPERATURE RESPONSE OF THE MAGNETIC HEAD

The magnetic head dissipates heat at a power that can be estimated as $P = RI^2$, and thus can increase its temperature when subject to large currents for prolonged periods of time. To avoid any interference with our measurements, such as heating of the sample, we mount the head on a large heat sink, that minimizes the increase on the temperature on the head.

We measure the temperature on the head surface for three different currents (350, 600 and 1000 mA) during 1 hour at time windows of 10 mn (Figure 8). The head just increases its temperature when subject to 1000 mA, by only $\sim 5°$ C, thus an insignificant increment. Furthermore, large currents are used to unfold proteins, and thus maintained for briefs periods of time ($\sim$ seconds). Prolonged currents are just maintained to observe folding dynamics at forces below 10 pN, where the head does not heat.

### VALIDATION OF THE CURRENT LAW

We check the ability of our current law to determine accurately the pulling force. In order to do so, we explore the folding/unfolding properties of protein L as a function of the force, and compare their dependency with that previously measured with conventional magnetic tweezers [9, 11]. Below 10 pN, protein L has a decreasing probability of populating the folded state, which can be well described by a sigmoid function as:

$$P_f = 1 - \frac{1}{1 + \exp{(8 - F)/0.93}}. \tag{21}$$

Above 10 pN, the unfolding rates of protein L can be well represented by a Bell-like exponential behavior as:

$$k_U(F) = k_0 \exp(F x^\dagger / kT), \tag{22}$$

with $k_0 = 0.0026$ s$^{-1}$, $x^\dagger = 0.44$ nm, and $kT = 4.11$ pNnm.

Figure 9 shows the folding probability (A) and unfolding rates (B) of protein L as measured with the magnetic head tweezers averaging data obtained at different values of $z$ and $I$. Black solid lines correspond to the expressions described above, which are fits to the data measured with conventional magnetic tweezers, validating our calibration of the magnetic head tweezers.

### VALIDATION OF THE CURRENT/DISTANCE DEPENDENCE OF THE PULLING FORCE USING TALIN DYNAMICS

We use talin to carry out a validation check of the dependence of the pulling force $F$ with the electric current $I$ and the distance $z$. In particular, we choose the R3 IVVI mutant talin domain, and design a polyprotein construct containing 8 repeats of the I91 titin domain, followed by the talin domain and the HaloTag (Fig. 10). The I91 domains work as the mechanical fingerprint.

The advantage of talin for our purpose here is that, being a force sensor, it has an exquisite force sensitivity, and changes in the pulling force of a fraction of a pN result in readily detectable changes in its folding probability—occupation of the folded state. Figure 10 shows the folding probability of the R3 IVVI domain measured with our magnetic tape head tweezers (black), and with the regular magnetic tweezers (red), to independently test this measurement. As it can be seen, talin transitions from fully folded to fully unfolded within less than 2 pN. At 8.7 pN, the population of the folded and unfolded states is the same. Just an increase in force of 0.3 pN (to 9 pN), drops this number to 30%. Hence, talin is an ideal system for testing the ability of our calibration to accurately predict the force over a large range of distances and currents.

Knowing that at 8.7 pN the talin hops between the folded and unfolded states with equal probability, we can predict the current $I$ needed at different distances $z$ to obtain this value of the force from Eq. 1 simply by rearranging terms:

$$I(z) = \frac{-9.462 + \sqrt{89.529 + 1.544F \arctan\left(\frac{12.5}{z}\right)\left[1 + \left(\frac{z}{12.5}\right)^2\right]}}{0.722 \arctan\left(\frac{12.5}{z}\right)}, \tag{23}$$

where $F = 8.7$ pN, and $z$ is expressed in $\mu$m and $I$ in mA. Hence, this equation can be used to predict the current $I$ needed to apply 8.7 pN at a particular distance $z$. Figure 3C demonstrates that it accurately predicts the experimental observations.

### KINETIC MODEL FOR THE NATIVE-MOLTEN GLOBULE FORMATION

We assume that the individual protein L domains reach the native conformation from the unfolded state through an intermediate state, which is characterized by low mechanical stability, and which we refer to as *molten globule.* As discussed in the main text, this is likely not a single well defined state, but rather an ensemble of collapsed state that contain part of the secondary structure present in the native conformation. Since the kinetics between this substates is very fast, and we cannot resolve them, we can gather them onto a single free energy basin, given the timescale separation between the weak events and the native ones. Thus, we assume the following kinetic model:

$$\mathrm{U} \underset{r_{\mathrm{MG}}}{\longrightarrow} \mathrm{MG} \underset{r_{\mathrm{N}}}{\longrightarrow} \mathrm{N} \tag{24}$$

where U stands for the unfolded state, MG for the molten globule, N the native state, and $r_{\mathrm{MG}}$ is the rate of formation of the molten globule from the unfolded domain, and $r_{\mathrm{N}}$ the rate of formation of the native state from the molten globule. Thus, we can write the following system of differential equations for $P_{\mathrm{U}}$, $P_{\mathrm{MG}}$ and $P_{\mathrm{N}}$, respectively the probability of observing the unfolded, molten globule and native state:

$$\frac{dP_{\mathrm{U}}}{dt} = -r_{\mathrm{MG}} P_{\mathrm{U}}, \tag{25}$$

$$\frac{dP_{\mathrm{MG}}}{dt} = r_{\mathrm{MG}} P_{\mathrm{U}} - r_N P_{\mathrm{MG}}, \tag{26}$$

$$\frac{dP_{\mathrm{N}}}{dt} = r_N P_{\mathrm{MG}}. \tag{27}$$

The population of the unfolded state is trivially $P_{\mathrm{U}} = P_{\mathrm{U}}^0 e^{-r_{\mathrm{MG}} t}$, being $P_{\mathrm{U}}^0$ the initial population of unfolded states, $P_{\mathrm{U}}^0 = 1$, in our case. This leaves us with the two coupled differential equations:

$$\frac{dP_{\mathrm{MG}}}{dt} = r_{\mathrm{MG}} e^{-r_{\mathrm{MG}} t} - r_{\mathrm{N}} P_{\mathrm{MG}}, \tag{28}$$

$$\frac{dP_{\mathrm{N}}}{dt} = r_{\mathrm{N}} P_{\mathrm{MG}}. \tag{29}$$

Considering the boundary conditions $P_{MG} = 0$ and $P_N(0) = 0$, we get:

$$\begin{aligned}
P_{\rm U}(t) &= e^{-r_{\rm MG}t}, \\
P_{\rm MG}(t) &= \frac{r_{\rm MG}}{r_{\rm MG} - r_{\rm N}} \left[e^{-r_{\rm N}t} - e^{r_{\rm MG}t}\right], \\
P_{\rm N}(t) &= \frac{r_{\rm MG}}{r_{\rm MG} - r_{\rm N}} \left(1 - e^{-r_{\rm N}t}\right) - \frac{r_{\rm MG} r_{\rm N}}{r_{\rm MG} - r_{\rm N}} t e^{-r_{\rm MG}t}.
\end{aligned} \tag{30}$$

Figure 11 shows the occupation of the unfolded (A), molten globule (B) and folded state (C) as a function of the quench time, measured in different 14 molecules, and a total of 546 events. Solid lines are global fits to the analytical expressions derived in Eqs. 30. The estimations for the rate of formation of the molten globule and folded states are: $r_{\rm MG} = 10.97 \pm 1.42$ s$^{-1}$, and $r_{\rm F} = 1.28 \pm 0.24$ s$^{-1}$. As it can be seen in the plots, the occupation of unfolded states does not decay to zero, and accordingly the occupation of native states does not saturate to one. This is due to the presence of some large steps associated to misfolded conformations with a contour length that encompass more than one protein L domain.

### DETAILED MATERIALS AND METHODS

#### Magnetic head tweezers setup

The presented custom-made setup is built on top of an inverted microscope (Olympus IX-71/Zeiss Axiovert S100) using a 63X oil-immersion objective (Zeiss/Olympus), mounted on a nanofocusing piezo actuator (P-725; Physik Instrumente) and a 1.6 X optivar lens. The fluid chamber was illuminated using a collimated cold white LED (Thor Labs). Images were acquired using a CMOS Ximea MQ013MG-ON camera. We applied force to paramagnetic Dynabeads$^{\rm TM}$M-270 beads (2.8 $\mu$m of diameter) through a magnetic head (BRUSH 902836) mounted on an articulating arm, and positioned with a micrometer with 1 $\mu$m resolution, and mounted on a heat sink. We control the current on the head by comparing the voltage drop across a precision resistance of 2 $\Omega$ with a computer-generated set-point, and feeding the error through a PID circuit (Stanford Research Systems SIM960). The PID output drives a 60 W audio amplifier (DROK 0 HI-FI Digital Stereo Power Amp), which drives directly the current to the head. The data acquisition and control of the piezo actuator was done using a multifunction DAQ card (NI USB-6289, National Instruments).

**Image processing**

Image processing was done by a custom-written software written in C++/Qt, fully available in the lab's website (http://fernandezlab.biology.columbia.edu/). Briefly, the z-position displacement of the bead was determined by first calculating the FFT of the image, computing the radial profile with a pixel addressing algorithm, and finally correlating the profiles from the magnetic and reference bead to a z-stack library calculated at the beginning of every experiment. With this procedure, we can reach frame rates up to 2 kHz, although the exact frequency depends on the size of the ROI in each experiment, determined by the relative position between the magnetic and reference beads.

**Fluid chamber preparation**

All magnetic head experiments were done in custom-made fluid chambers built by two sandwiched glass coverslips (Ted Pella), separated by a laser-cut pattern in parafilm. This pattern serves both to separate both glasses, and to create the fluidic wells where the buffers are added. The bottom surfaces where cleaned by sonication in a three-step protocol, involving first 30 min in 1% Hellmanex (Hellma), acetone, and ethanol (Sigma- Aldrich). Once dried with air, they are activated by air plasma during 20 min, and silanized by immersion for 15 min in a solution of (3-aminopropyl)- trimethoxysilane (Sigma-Aldrich) 0.1% v/v, in ethanol. After washing, the surfaces are cured at 100C for 1 h. Top glasses are cleaned by sonication in 1% Hellmanex for 30 min, and washed with ethanol. Next, they are immersed in a Repel Silane solution (Sigma-Aldrich) for 15 min to make them hydrophobic.

The custom-designed parafilm pattern is laser-cut and the bottom and top glasses sandwiched between it at 70C, to melt the parafilm. The fluid chambers are then functionalized in a three step protocol: 1) first, incubation for 1h with a solution of glutaraldehyde 1% v/v (Sigma-Adlrich) in PBS buffer 7.2 pH; 2) next, the chamber was filled with 0.025% w/v amine-terminated nonmagnetic polystyrene beads with diameter of 2.89 $\mu$m (Spherotech), diluted in PBS. After 10 min, the chamber is washed with 100 $\mu$L PBS buffer to eliminate the non adsorbed beads; 3) next, the chambers were reacted overnight with a solution of 10 $\mu$g/mL HaloTag amine (O4) ligand (Promega), diluted in PBS buffer. Finally the fluid chambers were blocked for 12 h with TRIS blocking buffer (20 mM Tris-HCL pH 7.4, 150 mM NaCl, 2mM $MgCl_2$, and 1% 2/v/ sulfhydryl blocked-BSA).

**Protein expression and purification**

Polyprotein constructs were engineered using a combination of *Bam*HI, *Bgl*II, and *Kpn*I restriction sites, as described previously. The protein constructs had eight repeats of protein L (B1 domain from *Peptostreptococcus magnus*, or eight I91 repeats followed by the mutant R3 IVVI talin domain, flanked by an N-terminal HaloTag enzyme (Promega) and a C-terminal AviTag for biotinylation. For purification purposes, a $His_6$-tag was also present before the AviTag. For expression and purification, the next steps were followed. BLR (DE3) or ERL competent cells containing the engineered pFN18A expression vector (Promega) were grown at 37 °C until $OD_{600nm} \approx 0.6 - 0.8$. The protein overexpression was induced with 1mM Isopropyl $\beta$-D-1-thiogalactopyranoside (IPTG, Sigma) overnight at 25 °C. Cells were resuspended in 50 mM sodium phophate buffer pH 7.0, 300 mM NaCl, 10% glycerol and 1mM Dithiothreitol (DTT, Sigma), and disrupted by French press. The proteins were purified from the lysate first with a Ni-NTA affinity purification resin, followed by size exclusion chromatography using a Superdex-200 HR column in 10 mM HEPES buffer pH 7.2, 150mM NaCl, 10% glycerol and 1mM EDTA. The purified proteins were then concentrated to 50-100 $\mu$M and biotinylated in 50 mM Bicine buffer pH 8.3, 10 mM magnesium acetate, 10 mM ATP, 100 $\mu$M biotin and 2.5 $\mu$g biotin ligase BirA enzyme (Avidity), at 40 °C for 4 hours.

**Single molecule measurements**

The protein was freshly diluted in HEPES buffer (Hepes 10 mM pH 7.2, NaCl 150 mM, EDTA 1 nM) containing 10 mM ascorbic acid (pH 7.3) to avoid oxidative damage, to a concentration of $\sim$ 50 nM, and left to adsorb on the surface for 30 min. Chambers where then washed with Hepes buffer, and streptaviding coated paramagnetic beads (Dynabeads$^{\text{TM}}$M-270, Invitrogen) were added to bind with the protein for $\sim$ 2 min. Next, the head was approached to a distance and a current value was supplied so that a force $\sim$ 4 pN was generated. Experiments begin with a z-stack library of the protein-tethered paramagnetic bead and a fixed reference bead. From then, the z-position displacements are calculated in real time using our image processing method, described previously [12].

**Single molecule data analysis**

The step sizes at unfolding and refolding transitions were measured from the individual trajectories at specific values of the current and the distance. Traces are previously smoothed with a Savitzky-Golay filter with $N = 101$, typically. Length histograms were obtained at each step, and the step size was determined by fitting double gaussian distribution to the histogram, and measuring the position of the consecutive histograms.

Protein L folding probability was measured from equilibrium trajectories by determining the residence time $t_{\mathrm{i}}$ of each folding state $i$ for a particular force value (measured at different currents and distances), and calculated as $P_{\mathrm{f}} = \sum_i it_{\mathrm{i}}/t_{\mathrm{t}}$, where the sum goes from $i = 0, \cdots, 8$ (being 8 the number of domains), and $t_{\mathrm{t}} = \sum_t t_{\mathrm{i}}$, the cumulative time at the particular force value. Talin folding probability, being a monomer, was determined by calculating the histogram of the length time-series of the smoothed trajectory $L(t)$, which shows a clear bimodal shape. Then, a double-gaussian fit was done, and the folding probability is calculated as the ratio between the area of both gaussians.

Unfolding rates for protein L were calculated by measuring the unfolding time for each domain $t_{\mathrm{i}}$, at a particular force value, measured at different currents and distances. Then, the unfolding rate is calculated as $r_{\mathrm{U}} = \left(\sum_i t_{\mathrm{i}}/N\right)^{-1}$, where $N$ is the number of unfolding events.

Molten globule experiments where done through a pulse protocol, with the following stages: 1) fingerprint pulse at 30 pN (or some other high force); 2) quench to 10 pN during 3 seconds; 3) quench pulse to 1 pN, during $\Delta t = 0.01$, 0.02, 0.05, 0.1, 0.25, 0.5, 1, 5, and 10 seconds; 4) probe pulse at 10 pN during 100 seconds; 5) refolding pulse at 4 pN during 100 seconds. The unfolding time histogram was generated by measuring the unfolding time $t_{\mathrm{U}}$ of any event observed during pulse (4), and calculating the histogram of the $\ln(t_{\mathrm{U}})$. Steps sizes of this events, where also monitored and plot in two different histograms; all step sizes resulting in less than 2.05 seconds where accounted as molten globule events, and all step sizes with unfolding times higher than 2.05 seconds as native domains. Finally, the probability of molten globules $P_{\mathrm{MG}}$ and native domains $P_{\mathrm{N}}$ as a function of the quench time $\Delta t$ was calculated by measuring the number of molten globules $N_{\mathrm{MG}}$ and folded domains $N_{\mathrm{N}}$, only for those events with step sizes between 9 and 11.5 nm (standard deviation of the gaussian fit at 10 pN), and computed as $p_{\mathrm{MG}} = N_{\mathrm{MG}}/N$, $p_{\mathrm{N}} = N_{\mathrm{F}}/N$, where $N$ was the number of domains unfolded in the fingerprint pulse (typically $N = 8$, but we also accepted pulses with $N = 7$, where 1 domain was lost due to oxidative damage).

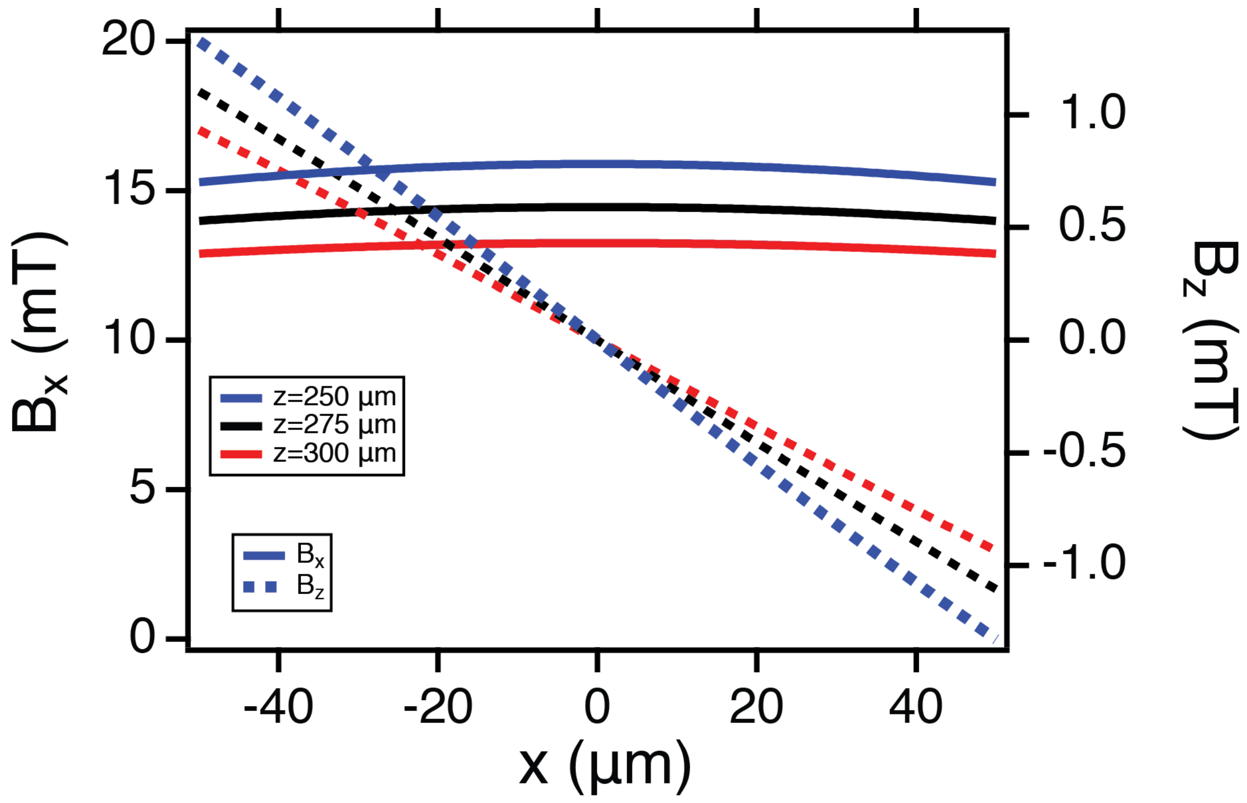


FIG. 1. Karlqvist components in $x$ (solid) and $z$ (dotted) for three different working distances.

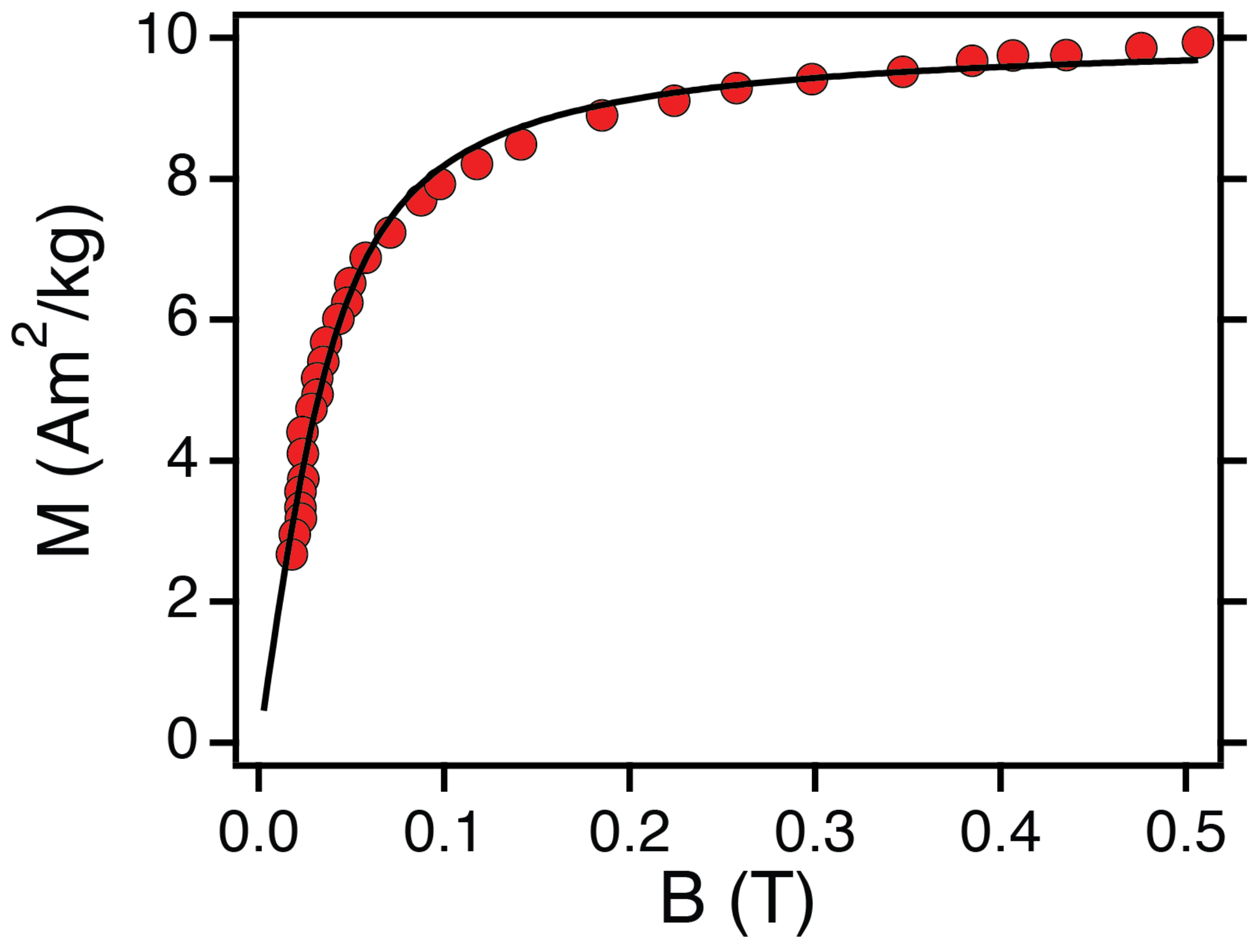


FIG. 2. Magnetic response of Dynabeads$^{TM}$M-270, fit to a Langevin magnetization model (data taken from [10]).

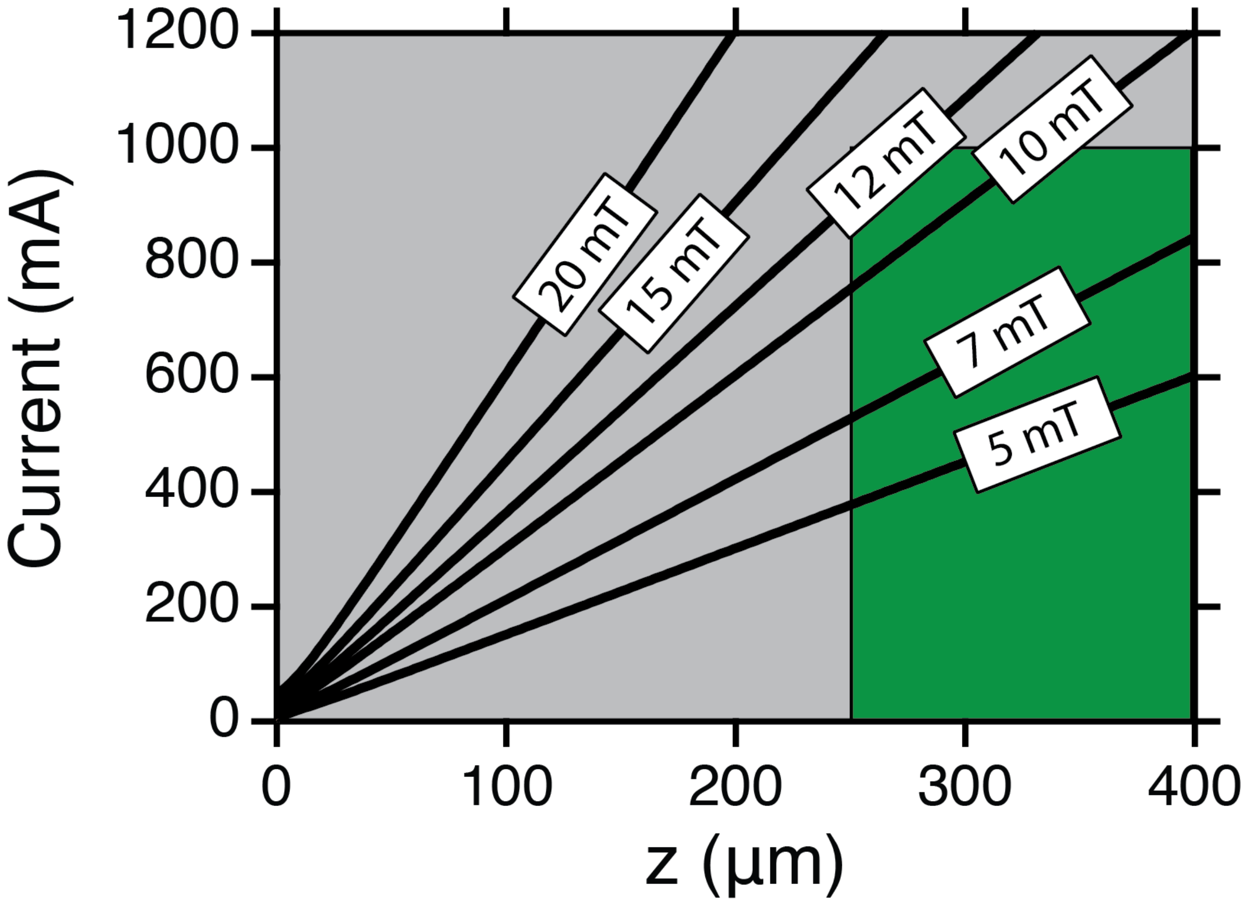


FIG. 3. Current versus distance relation for different values of the field. Green box shows the working region of our setup ($250 \mu m < z < 400 \mu$m; and $0 < I < 1$ A).

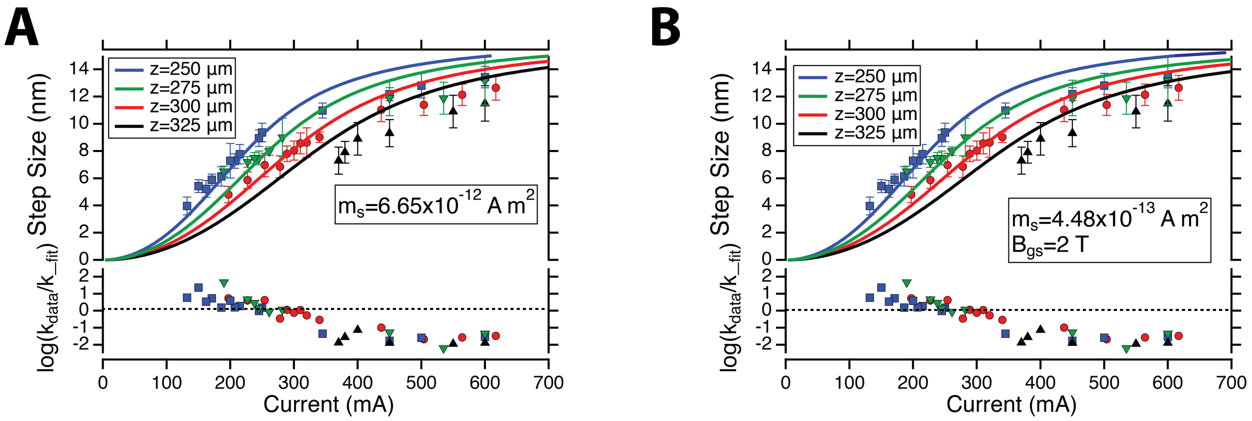


FIG. 4. Global fit of the step sizes to the current law developed from Langevin approximation of the magnetization, leaving one single parameter ($m_{\mathrm{s}}$) free, or two ($m_{\mathrm{s}}$, and $H_{\mathrm{gs}}$).

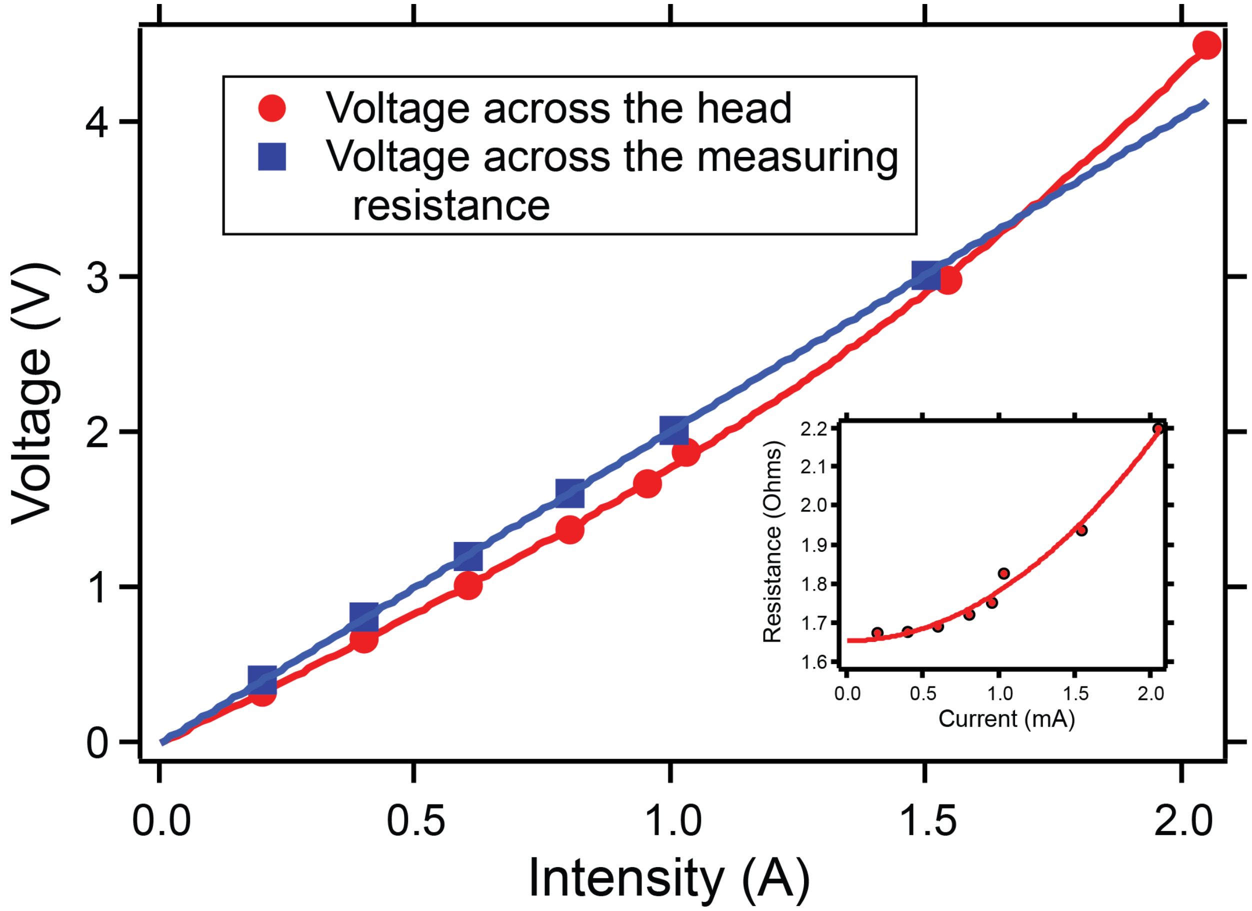


FIG. 5. Measurement of the voltage across the head (red) and across a measuring resistance of 2 Ω (blue) at different fixed currents. The head heats at large currents resulting in a non-constant value of the resistance.

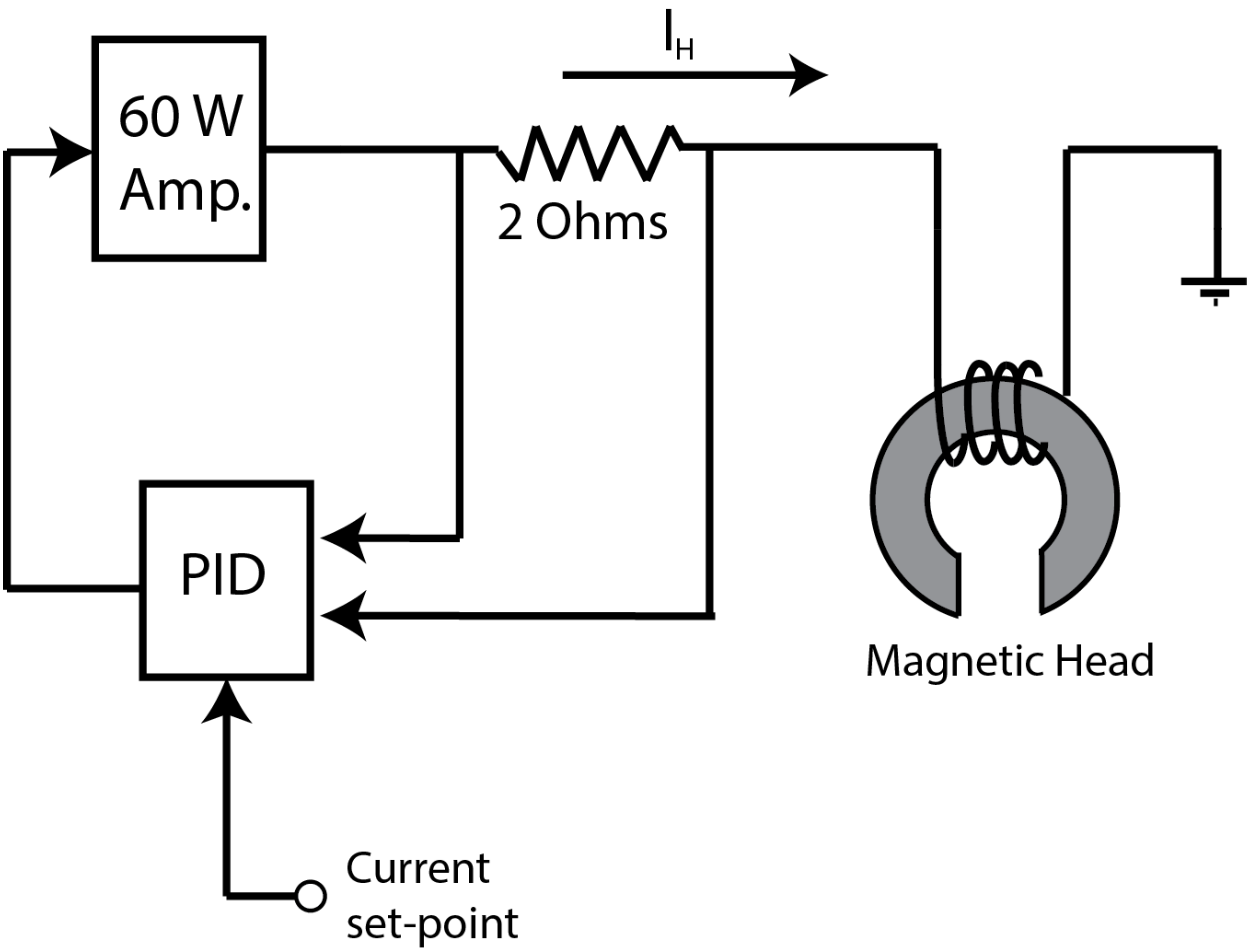


FIG. 6. Schematic diagram of the current control circuit for the magnetic head. We measure the current across the head ($I_H$) by measuring the voltage drop across a high power 2 $\Omega$ precision resistor. The voltage drop is compared with a computer generated current set-point and the error is fed through a PID circuit (Stanford Research Systems SIM960) whose output drives a 60 W audio amplifier (DROK 0 HI-FI Digital Stereo Power Amp). The output of the power amplifier directly drives current through the head in response to the value of the set-point. The bandwidth of the circuit is $\sim$ 20 kHz.

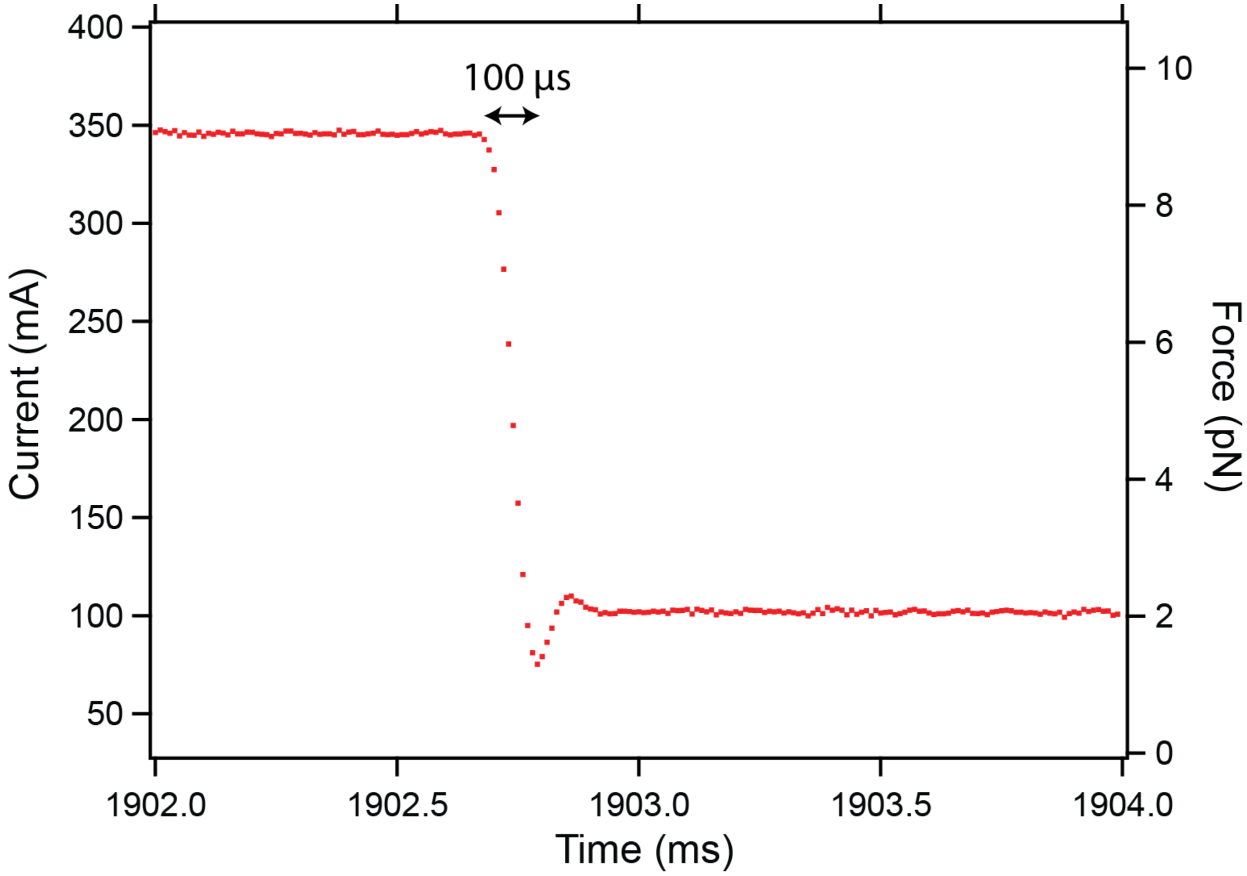


FIG. 7. Direct measurement of a step in current from 345 mA to 102 mA (left axis), which at a distance of 300 $\mu$m, corresponds to a change in force from 9 pN to 2 pN. The current is sampled at 100 kHz. With our PID controller and the 60 W power amplifier, the current can change within 100 $\mu$s, as it can be seen in the figure. The effect of the PID on the relaxation of the electric current can be observed at this very short timescale.

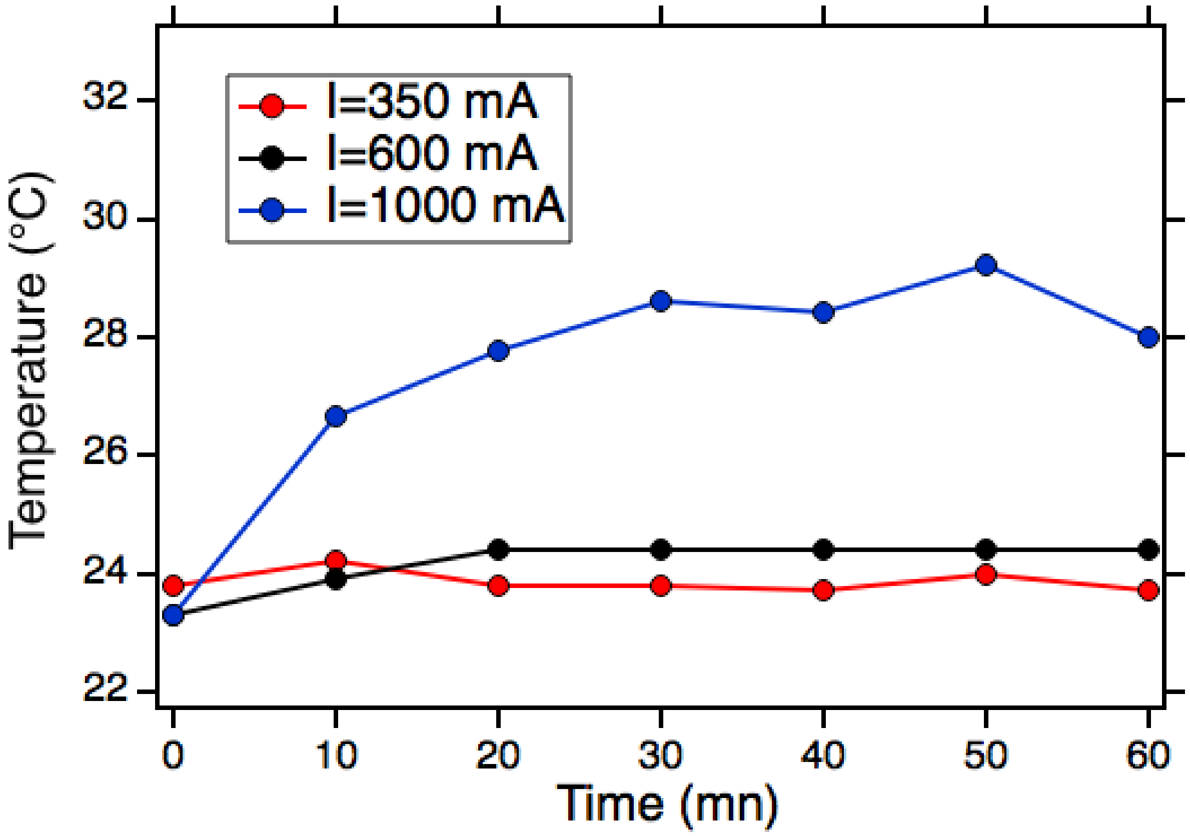


FIG. 8. Temperature on the head surface at different currents and time windows.

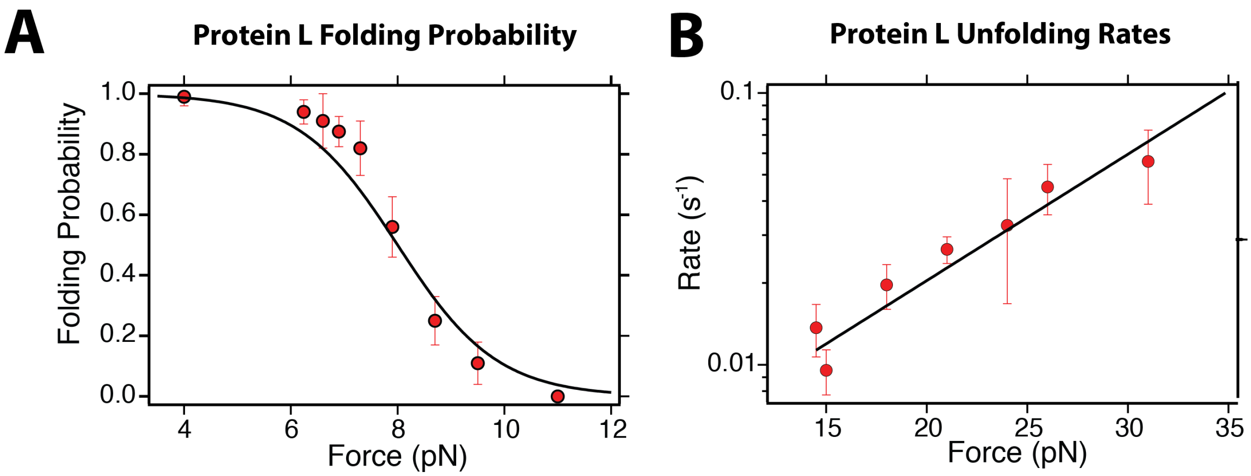


FIG. 9. Folding probability (A) and unfolding rates (B) measured with magnetic head tweezers, and with conventional magnetic tweezers (solid lines). Data measured with 5 molecules, at 4 different distances (300, 275, 265, and 250 $\mu$m).

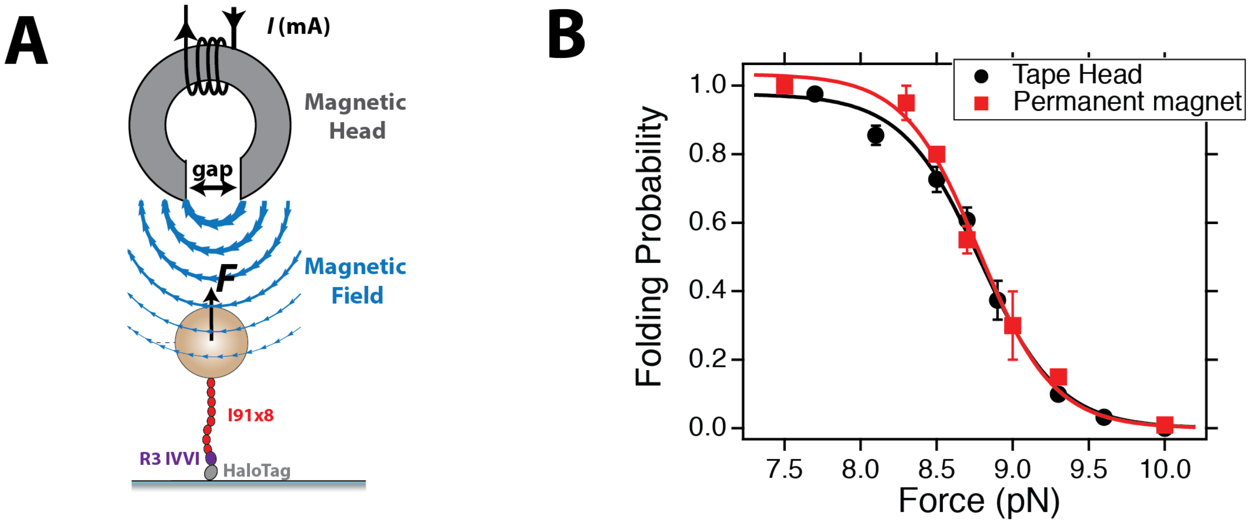


FIG. 10. (A) Schematics of the experimental setup for talin experiments, highlighting the molecular construct, which contains eight I91 domains (red) and the talin R3 IVVI mutant domain (purple). (B) Folding probability of talin R3 IVVI mutant domain measured with the tape head tweezers (black) and permanent magnets (red). Data measured over $> 3$ molecules.

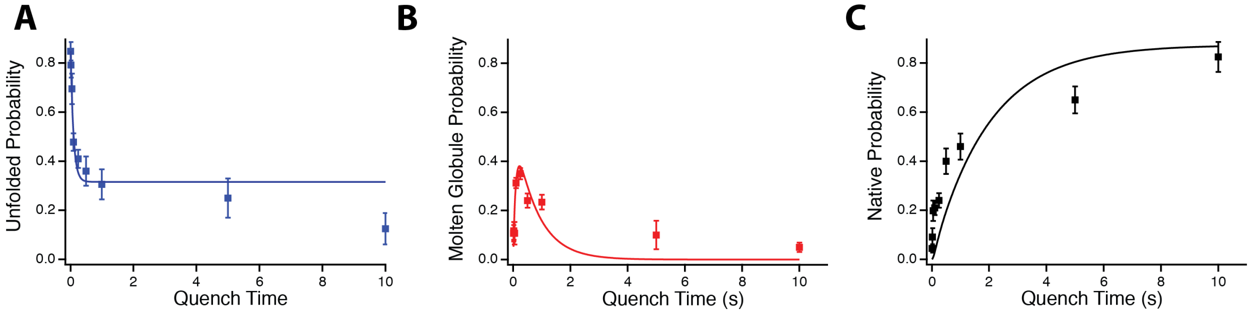


FIG. 11. Occupation of the unfolded (A), molten globule (B) and native (C) states as a function of the quench time at 1 pN. Solid lines are global fits to Eqs. 30 with the rate of formation of the molten globule $r_{\mathrm{MG}}$ and native state $r_{\mathrm{N}}$ as fitting parameters, which yields $r_{\mathrm{MG}} = 10.97 \pm 1.42\ \mathrm{s}^{-1}$, and $r_{\mathrm{F}} = 1.28 \pm 0.24\ \mathrm{s}^{-1}$. Data collected with 14 molecules, and a total of 546 events.

TABLE I. Magnetic properties of Dynabeads™M-270. CV is the standard deviation in the bead diameter, as a percentage of its mean diameter; $\chi$ is the initial susceptibility (dimensionless); and $M_s$ is the mass saturation magnetization of the sample.

| Bead | Diameter ($\mu$m) | CV (%) | $\rho$ (kg/m$^3$) | $\chi$ | $M_s$ (Am$^2$/kg) |
|---|---|---|---|---|---|
| M-270 | 2.83 | 1.4 | 1400 | 0.756 | 10.8 |

TABLE II. Technical Specifications the magnetic head Brush Industries, 902836.

| | |
|---|---|
| Core Material | Ceramic-coated Hypermium |
| Gap Length | 0.001 inch (25 $\mu$m) |
| Inductance | 0.4 mH |
| D.C.R. | 1.4 to 1.8 $\Omega$ |
| Saturation Current | 600 mA (0 - P); 1.2 A in DC |
| Maximum Gap Field | 500 mT |

TABLE III. Probability of occupation of the folded state after a 250 ms long quench. Error is the SEM. Probabilities calculated over 43, 50, 64, and 72 folding pulses, respectively, and a total of 8 molecules.

| Force (pN) | 0.5 | 1.0 | 2.0 | 4.0 |
|---|---|---|---|---|
| $P_f$ | 0.25±0.03 | 0.25 ± 0.05 | 0.28 ± 0.02 | 0.49± 0.03 |
